# In vitro coinfection by influenza A virus and respiratory syncytial virus generates hybrid viral particles with altered structure and tropism

**DOI:** 10.1101/2021.08.16.456460

**Authors:** Joanne Haney, Swetha Vijayakrishnan, James Streetley, Kieran Dee, Daniel Max Goldfarb, Mairi Clarke, Margaret Mullin, Stephen D Carter, David Bhella, Pablo R Murcia

## Abstract

Interactions between respiratory viruses impact viral transmission dynamics and clinical outcomes. To identify and characterize virus-virus interactions at the cellular level, we coinfected human lung cells with influenza A virus (IAV) and respiratory syncytial virus (RSV). Super-resolution microscopy, live-cell imaging, scanning electron microscopy, and cryo-electron tomography revealed extracellular and membrane-associated filamentous structures consistent with hybrid viral particles (HVPs). We show that HVPs harbor surface glycoproteins and ribonucleoproteins of IAV and RSV, and use the RSV fusion glycoprotein to evade anti-IAV neutralising antibodies and to infect and spread among cells lacking IAV receptors. Finally, we show evidence of IAV and RSV coinfection within cells of the bronchial epithelium, with viral proteins from both viruses co-localising at the apical surface. Our observations have profound implications for infection biology as they define a previously unknown interaction between respiratory viruses that might affect virus pathogenesis by expanding virus tropism and facilitating immune evasion.

## Main Text

Viruses are intracellular pathogens whose replication relies on the infection of a restricted subset of cell types, a property known as tropism. As multiple viruses exhibit the same tropism, cells and tissues are susceptible to infection by a community of taxonomically diverse viruses. From this perspective, a tissue or body compartment constitutes an ecological niche in which members of a virus community co-exist. Coinfections by more than one virus represent between ∼10-30% of all respiratory viral infections and are common among children (*1, 2*). The clinical impact of viral coinfections is unclear: while some studies indicate that coinfections do not alter the outcome of disease (*3-5*), others report increased incidence of viral pneumonia (*6, 7*). At the cellular level, the underlying interactions that determine the outcome of coinfections are unclear. Direct interactions between viruses within coinfected cells can result in changes to viral progenies, including -but not limited to-pseudotyping (incorporation of surface proteins from a different virus) (*8-10*) or genomic rearrangements, which may generate novel strains with pandemic potential such as SARS-CoV-2 and pandemic influenza A viruses (*11, 12*). Here, we examined interactions between two commonly co-circulating viruses of clinical importance: influenza A virus (IAV) and respiratory syncytial virus (RSV). IAV causes over five million hospitalizations each year (*13*) and RSV is the leading cause of acute lower respiratory tract infection in children under five years of age (*14, 15*).

To study virus-virus interactions during confection, we infected a cell line derived from human lung (A549) (*16*), with a mixed inoculum of IAV and RSV, or individual viruses as controls. Infections were performed at high multiplicity of infection (MOI) to facilitate coinfection and recapitulate high MOIs produced in advanced stages of infection (when IAV and RSV foci may come into contact). We compared viral replication in single infections and coinfections. Single IAV infections resulted in robust replication, peaking between 24-48 hours post-infection (hpi). In coinfections, IAV replication was similar to that observed in coinfection or at some timepoints marginally increased (Fig. 1a, top left). As RSV titres peaked in single infections at a later timepoint than IAV (72 hpi, Fig. 1a, top right), further infections were performed to determine optimal conditions for coinfections by using a reduced IAV MOI (ten-fold reduction relative to RSV). The reduced IAV input did not affect IAV replication, which still replicated to higher titres in coinfections (Fig. 1a, bottom left). In contrast, RSV titres were significantly reduced, by over 400-fold at 72 hpi, in the presence of IAV both at equivalent MOI (Fig. 1a, top right) and when IAV MOI was reduced relative to RSV (Fig. 1a, bottom right). These results are consistent with published studies that show that RSV is adversely impacted by coinfection, while IAV replication is not (*17, 18*).

**Figure 1.**
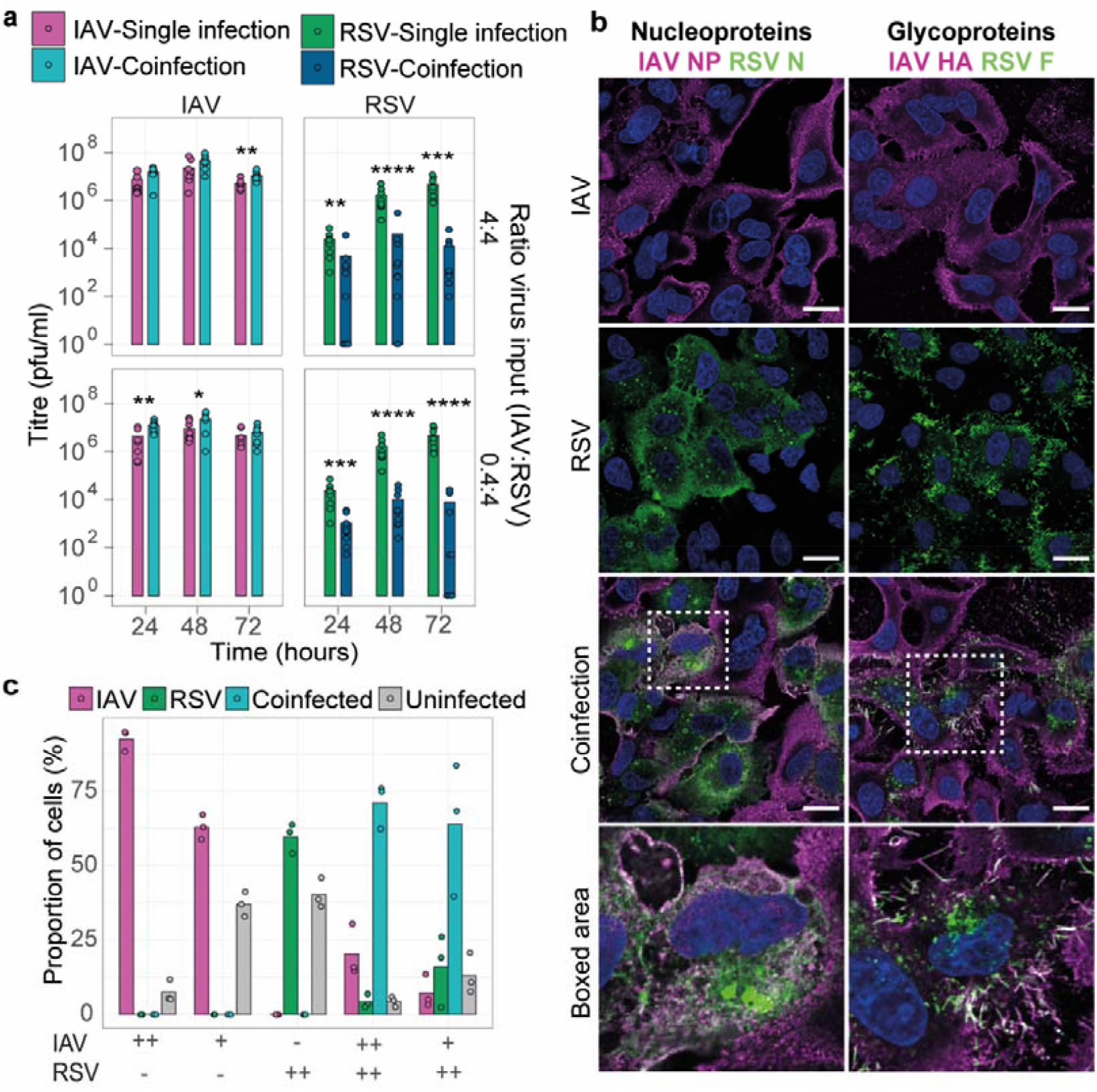
Replication kinetics and levels of cellular infection are affected in IAV and RSV coinfections. (a) Left panel, IAV replication kinetics in single infection (magenta bars) and mixed infection with RSV (teal bars). Right panel, RSV replication kinetics in single infection (green bars) and mixed infection with IAV (blue bars). Top row, infections carried out at equivalent MOI (4:4), bottom row, IAV input 10-fold less than RSV input (0.4:4). Error bars represent standard error, significance determined by Mann-Whitney test, * p<0.05, ** p<0.01, *** p<0.001, **** p<0.0001, non-significance not indicated. (b) Single or coinfected A549 cells were fixed at 24 hpi and stained by immunofluorescence for IAV nucleoprotein (magenta) and RSV nucleoprotein (green) (left column) and IAV haemagglutinin (magenta) and RSV fusion glycoprotein (green) (right column). Nuclei are stained with DAPI (blue). White boxes indicate regions in labelled boxed area, scale bar indicates 20 µm. (c) Percentage of cells infected with IAV-only (magenta), RSV-only (green), coinfected (teal) or uninfected (grey) in single virus infection or mixed at 24 hpi. Panel below plot represents the infection conditions used: - for no virus, ++ for MOI 4 and + for MOI 0.4.

To assess if coinfection affected localization of key viral proteins or the proportion of infected cells, single and coinfections were performed as described above and cells were fixed at 24 hpi. Immunofluorescence microscopy revealed that cells infected with either RSV or IAV displayed features typical of the replicative cycle of each virus, such as cytoplasmic inclusion bodies containing RSV nucleoprotein (N), or diffuse IAV nucleoprotein (NP) staining within the cytoplasm (Fig. 1b, left column). Similar features were observed in coinfected cells (Fig. 1b, boxed area) suggesting that coinfection did not affect the intracellular localization of viral proteins. We counted the number of infected cells in single infections alongside the number of single infected and coinfected cells in the coinfection condition. We observed a high proportion of coinfected cells at 24 hpi when cells were infected with equivalent amounts of virus as well as when IAV input MOI was reduced 10-fold relative to RSV (Fig. 1c), suggesting that superinfection exclusion does not prevent coinfection. Paradoxically, a higher number of RSV-positive cells were detected in coinfections than in RSV single-infected samples, contrasting with the observed reduction in RSV titre in coinfections.

IAV and RSV are enveloped viruses that target lipid rafts for assembly and budding (*19, 20*). We therefore stained cells for the major glycoproteins of IAV and RSV: hemagglutinin (HA) and RSV fusion (F), respectively (Fig. 1b, right column) and examined their localisation at plasma membranes. IAV HA diffusely coated the plasma membrane of infected cells in both single virus infections and coinfections, while RSV F was localized to regions containing filamentous structures extending from the cell surface, corresponding to budding RSV virions. In coinfected cells, HA was not excluded from RSV budding sites (Fig. 1b, right column, boxed area). We thus hypothesized that mixing of glycoproteins within these regions may result in incorporation of both IAV and RSV components to budding viral particles. To test this, we applied super-resolution confocal microscopy to image virions immunolabelled for HA and F at high resolution. We observed dual-positive fluorescence suggesting the presence of budding viral filaments possessing both IAV and RSV glycoproteins during coinfection (Fig. 2a, b). Strikingly, minimal colocalization between the two glycoproteins was observed. Instead, glycoproteins were incorporated in distinct regions along the filament, with most dual-positive filaments positive for IAV HA at the distal end (Extended Data Fig. 1). 56±9.1% (mean ± SD) of coinfected cells displayed cell-associated dual-positive filaments. Many of these cells showed extensive production of dual-positive filaments, with no obvious indication of cell damage (Fig. 2c, d). We also observed dual-positive filaments distant from coinfected cells, suggesting they remained intact upon budding (Fig. 2b). Nested among dual-positive filaments (both cell-associated and released), we identified filaments corresponding to IAV and RSV virions (identified by HA or F only staining) (Fig. 2a, b, Extended Data Fig. 1). This indicates that regions in which dual-positive filaments form may contain both IAV and RSV budding sites.

**Figure 2.**
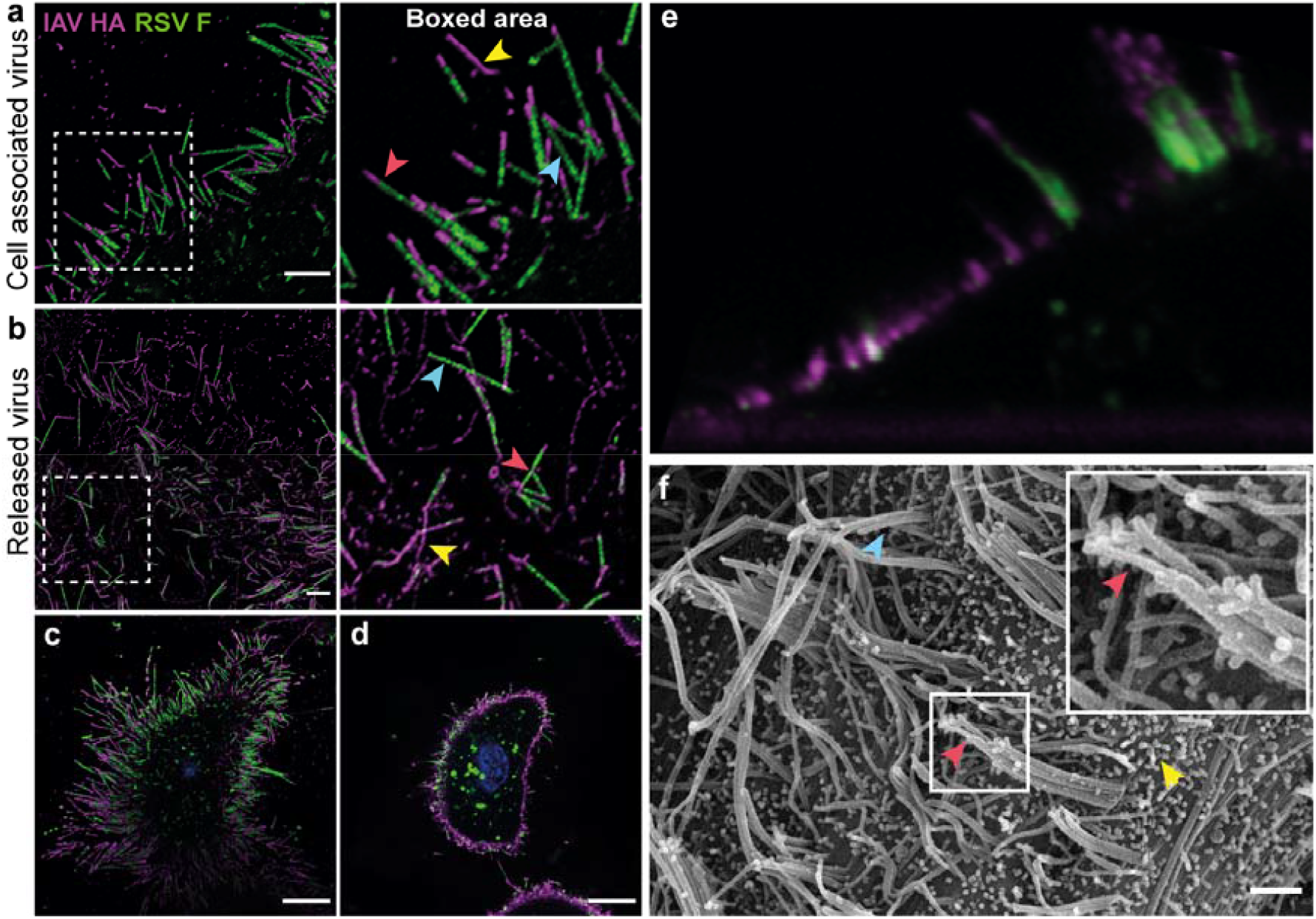
Coinfected cells display budding viral particles incorporate elements of both IAV and RSV. Cell associated viral filaments positive for IAV HA (magenta on merged image) and RSV F (green on merged image) that are cell associated (a) or released (b) from coinfected cells, imaged by super-resolution microscopy. Magnified images of white boxed regions indicate examples of IAV filaments (yellow arrows), RSV filaments (blue arrows) and dual-positive filaments (red arrows). Scale bars indicate 500 nm. (c and d) Coinfected cell exhibiting extensive dual-positive filament formation. Cell imaged by confocal microscopy at two Z-positions: basal side adhered to coverslip (c) and through the centre of the cell (d). Scale bars indicate 10 µm. (e) 2D z-plane image reconstructed from a Z-stack of a live cell stained for IAV HA and RSV F, imaged by super-resolution confocal microscopy at 100 nm intervals through the entire cell (16.3 µm). (f) Scanning electron micrograph of coinfected cell surface. IAV and RSV particles are indicated by yellow and blue arrows respectively. Red arrow and magnified region highlight particles that contain structural features of both IAV and RSV. Scale bar indicates 1 µm.

We performed microscopy on live cells to visualize the native organization of the filaments budding from coinfected cells. Consistent with our previous observations, we detected bundles of filaments protruding from the cell surface that incorporated F along the length of the filament with HA localized at the distal end (Fig. 2e). To obtain better resolution of these budding viruses, we used scanning electron microscopy (SEM). IAV single infection revealed a pleomorphic population of membrane-associated structures consistent with spherical, bacilliform and filamentous particles, whereas RSV single infection resulted in dense clusters of budding RSV filaments (Extended Data Fig. 2). In coinfected cells we identified RSV–like filaments with structures branching from the distal ends resembling smaller bacilliform IAV-like virions (Fig. 2f).

To determine the structural details of IAV/RSV filamentous structures budding from coinfected cells, we performed cryo-electron tomography (cryo-ET) and observed structures consistent with hybrid viral particles containing elements of both IAV and RSV. We repeatedly observed two classes of hybrid particles with some virions exhibiting features of both classes: in 11 cryo-tomograms we observed true hybrid virus particles (HVPs, Fig. 3), and in 16 cryo-tomograms we observed pseudotyped viruses (PVs, Extended data Fig. 3). Pleomorphic IAV particles and RSV filaments were also identified (Extended data file 1, Extended Data Movies 1-2).

**Figure 3:**
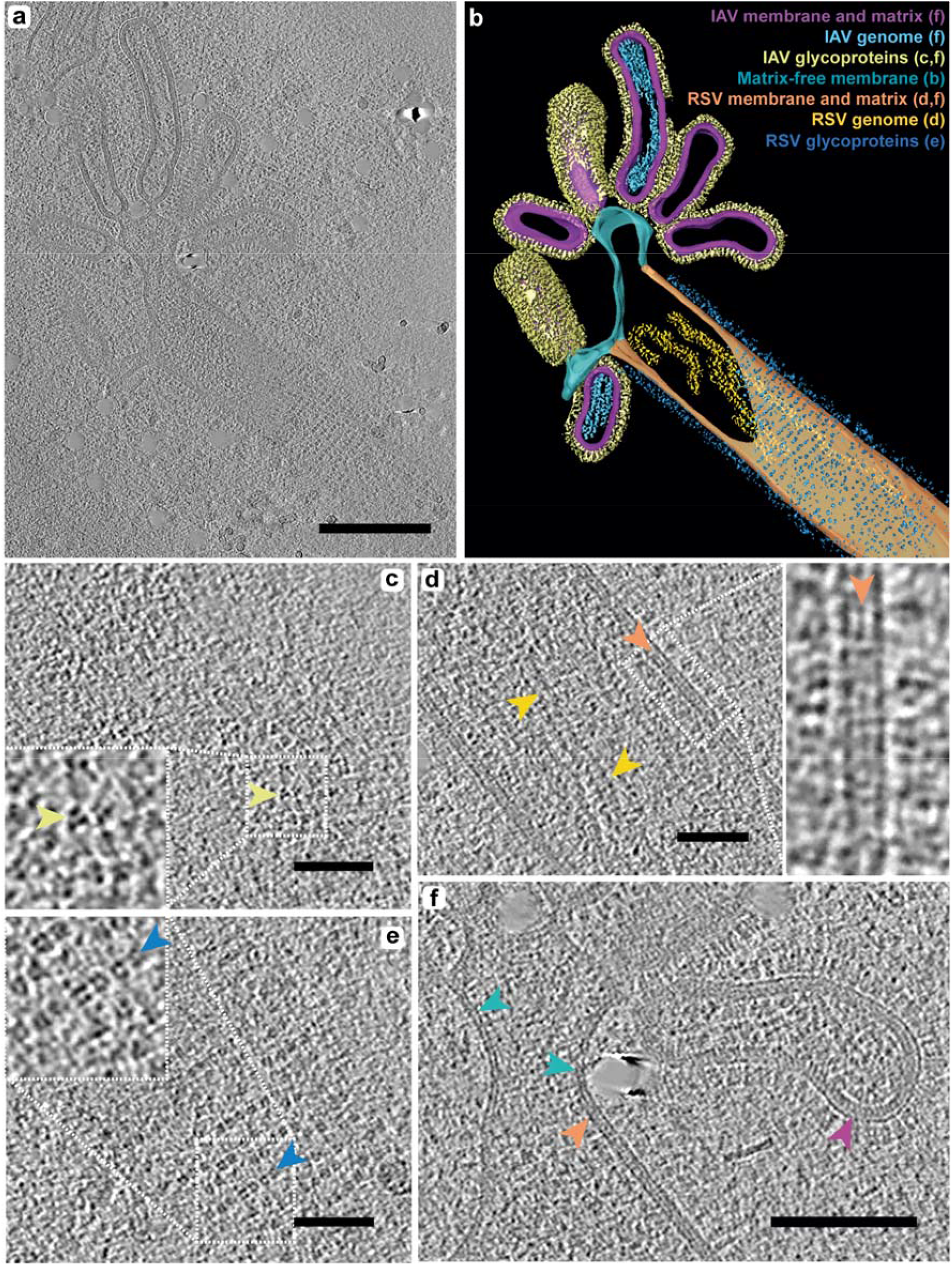
Coinfected cells produce hybrid viral particles that contain IAV- and RSV-like structural regions. Viral particles produced from coinfected A549 cells were frozen at 24 hpi and imaged by cryo-ET. (a) Cryo-tomogram showing an example of a hybrid viral particle. Particles consisted of an RSV-like region, extending to multiple IAV-like protrusion from the distal end of the filament. Scale bar indicates 200 nm. (b) Manual segmentation of the cryo-tomogram shown in (a) highlighting examples key structural features (not all features have been segmented). Coloured labels on figure match-coloured segmented volumes and letters denote figure panels in which detailed images of features are shown. (c) The surface of IAV region shows the ordering of IAV glycoproteins (yellow arrow). Magnified insert shows density associated with HA trimers in a characteristic triangular arrangement. (d) Cross-section through the RSV region shows RNPs, indicated by gold arrows. The inset image shows magnified region highlighted by white dashed box, showing the RSV viral envelope associated with the matrix layer (orange arrow). (e) The surface of RSV region shows the characteristic helical arrangement of glycoproteins into rows (blue arrows). The magnified insert shows circular density corresponding to RSV glycoproteins. Scale bars in panels (c-e) indicate 50 nm. (F) IAV- and RSV-like regions were joined by a continuous membrane bilayer. Coloured arrows indicate presence or absence of matrix layer. Membrane in IAV and RSV region was clearly associated a with matrix layer, indicated by orange and magenta arrows for RSV and IAV respectively, whilst matrix appeared absent in the adjoining regions (blue arrow). Scale bar indicates 100 nm.

HVPs exhibited distinct IAV-like and RSV-like structural regions: the wider part of the filament resembled RSV and extended continuously towards the distal end to one or more narrow, IAV-like regions (Fig. 3a-b, Extended data Fig. 4, Movies 1 and 2). Density consistent with both IAV and RSV ribonucleoproteins (RNPs) could be visualized and seemed confined within their respective structural regions of the filaments (*21, 22*) (Fig. 3a, b and d). Glycoproteins consistent in size, shape and density with IAV or RSV glycoproteins decorated the respective IAV or RSV-like regions of the particles (Fig. 3c and e). In some virions, the junction between IAV and RSV regions had a clear lumen (Extended Data Fig. 4c, Movie 2), whilst other particles exhibited a much narrower join, with pinching off membranes between IAV and RSV areas (Fig. 3f, Movie 1, Extended Data Fig. 4a). Matrix layers were observed in IAV and RSV regions but were absent from the membrane bridging the two regions (Fig. 3f).

We also identified virions consistent with RSV filaments containing (containing RSV RNPs) that had a different glycoprotein ordering than expected (Extended data Fig. 3). The glycoproteins differed in shape when compared to those present on RSV virions and had a triangular arrangement which resembled IAV glycoproteins (Extended data fig. 3c), rather than ring shaped density more consistent with RSV glycoproteins (Extended data fig. 3e). They also lacked the long-range helical ordering typical of RSV glycoproteins (Extended data fig.3c and e) (*23*). We designated these structures *pseudotyped viruses* (Extended Data Fig. 3a, c and d, Movie 3). Some HVPs with distinct IAV- and RSV-like structural regions also exhibited pseudotyping within the RSV region (Extended Data Fig. 4a-b). Notably, we did not observe IAV virions or IAV-like regions in hybrid particles that were pseudotyped with RSV glycoproteins. To quantify differences in glycoprotein arrangement, we determined the inter-spike distance on IAV, RSV and PVs. RSV exhibited a mean (+/- SD) inter-spike spacing of 12.9 (+/-2.32) nm, while IAV had a spacing of 8.71 (+/-1.18) nm. PVs had an average spike distance ranging from 8.31-9.56 nm, similar to IAV but significantly different to RSV (p<0.0001, unpaired t-test) (Extended Data Fig. 5). These results confirm that coinfections can result in the formation of two classes of hybrid viral particles structurally distinct from either parental virus.

As surface glycoproteins determine antigenicity and tropism, and HVPs incorporate glycoproteins of both IAV and RSV, we hypothesized that HVPs would display altered antigenicity. To test this, we first compared the neutralization efficiency of anti-IAV hemagglutinin (HA) antibodies against viruses collected from cells infected with IAV alone, or coinfected with IAV and RSV. As RSV is predominantly cell-associated, we performed neutralization assays using supernatant and cell-associated fractions (see methods). Viruses were also back-titrated to determine infectious titre of the inoculum (Extended data Fig. 6) Viruses collected from coinfected cells displayed reduced IAV neutralization compared to those collected from single IAV infections (Fig 4a). While the observed differences were not statistically significant, the decrease in neutralization efficiency was more marked in the cell-associated fraction of the coinfected cells (Fig. 4a) where only 33% (+/- 27%) (mean [+/- SD]) of IAV was neutralized, suggesting that two-thirds of IAV within this fraction was able to evade antibody-mediated neutralization (Fig. 4a). In an analogous experiment, we compared the neutralization efficiency of anti-RSV F antibodies (Palivizumab) against viruses collected from cells infected with RSV alone or coinfected. RSV harvested from both single and mixed infections was efficiently neutralized (Fig. 4b). This suggests that, in contrast to IAV, RSV cannot utilise IAV glycoproteins to facilitate viral entry. Further, in the context of PVs, RSV infectivity may be determined by the ratio of incorporated IAV and RSV glycoproteins, where fully pseudotyped RSV filaments may be non-infectious, as they are unable to utilise IAV HA.

**Figure 4:**
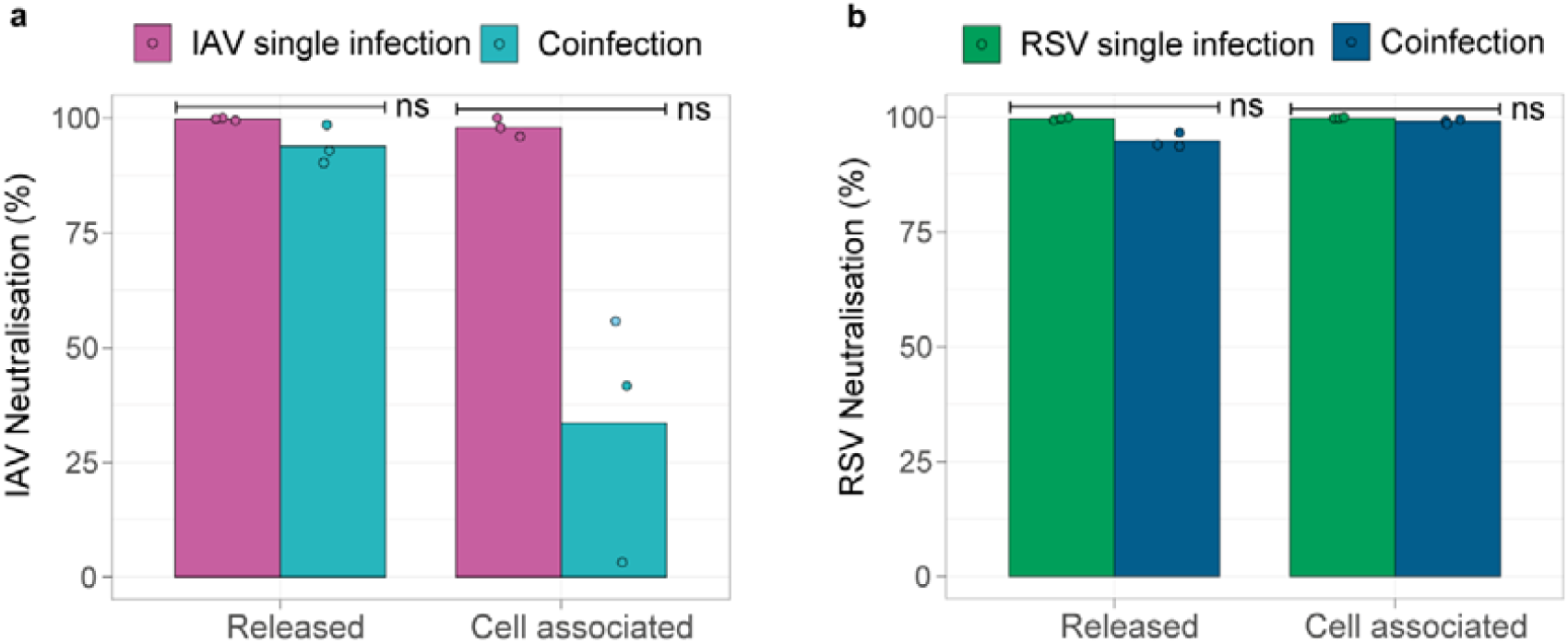
Virus produced from coinfection facilitates evasion of neutralising antibodies by IAV, but not by RSV. Virus harvested from coinfection or single infections was pre-incubated with serum targeting IAV HA, RSV F or a serum-free control, and then used to infect A549 cells. Infections were fixed and immunostained at 12 hpi and the number of IAV infected cells was quantified using an automated image-based cell counter. (a) Neutralisation of IAV by polyclonal antisera targeting IAV HA when virus was harvested from the supernatant or cell pellet fractions of a single IAV infection (magenta bars) or a coinfection (teal bars). Neutralisation was calculated as a percentage of IAV infected cells in the wells containing serum compared to matched serum-free controls. Statistical significance calculated by Mann Whitney test, ns indicates p>0.05. (b) Neutralisation of RSV by Palivizumab (targeting RSV F) when virus was harvested from the supernatant or cell pellet fractions of a single RSV infection (green bars) or a coinfection (dark blue bars). Neutralisation calculated as a percentage of RSV infected cells in the wells containing serum compared to matched serum-free controls. Data was collected from three independent experiments and statistical significance calculated by Mann Whitney test, ns indicates p>0.05.

To determine if the incorporation of RSV glycoproteins could result in an expansion of IAV receptor tropism, we treated cells with neuraminidase (NA) to remove sialic acids (the cellular receptor for IAV) (Extended Data Fig. 7a). Sialic acid removal was confirmed by lectin staining (Extended Data Fig. 7b). Virus was harvested from IAV or RSV single infections or coinfections as described for neutralisation assays and subsequently used to inoculate IAV-receptor-deficient (NA-treated) or control cells (Fig. 5a), as well as being back-titrated to determine the IAV titre in the inoculum (Extended data Fig. 7c). Cells were fixed at 12 hpi, immunostained for IAV NP or RSV N and infected cells quantified. IAV entry was blocked in NA-treated cells when inoculated with the released virus of single IAV-infected cells, whereas entry of cell-associated IAV collected from single IAV infections was reduced by 90% (Fig. 5a). When NA-treated cells were inoculated with released or cell-associated virus harvested from mixed infections, IAV entry was significantly increased compared to IAV-only infection (Fig. 5a). The increase in IAV entry was higher in the cell-associated fraction, and IAV entry in receptor-deficient cells was restored to 63% (+/- 10%) (mean [+/- SD]) of the level of control cells (Fig. 5a). As RSV does not use sialic acids as receptors, no differences in RSV entry between NA-treated or control cells were detected (Extended Data Fig. 7d). To determine if extracellular association between free IAV and RSV virions was contributing to IAV entry in NA-treated cells, IAV and RSV were mixed and incubated prior to infecting NA-treated or control cells. No significant differences were observed in IAV or RSV entry into receptor-deficient cells compared to control cells when the viruses had been pre-mixed (Extended data Fig. 7e), compared to each virus alone, suggesting that IAV and RSV do not associate upon extracellular mixing or that association does not impact viral entry. Overall, our findings suggest that the increase in IAV entry to sialic acid-deficient cells was a result of hybrid particle formation and indicate that HVPs represent a subpopulation of infectious virus particles with expanded receptor tropism. To determine whether the expansion of IAV tropism was mediated by the RSV F glycoprotein, virus harvested from mixed infections was incubated with Palivizumab. We observed a significant reduction in entry of both released and cell-associated IAV into NA-treated cells in the presence of Palivizumab (Fig. 5b-c), suggesting that RSV F facilitates IAV infection via hybrid particles that enable IAV to enter cells that would otherwise be refractory to infection. We measured IAV entry into untreated control cells (i.e. cells with normal expression of sialic acids) in the presence or absence of Palivizumab. We observed that in the presence of Palivizumab, entry of IAV harvested from the cell associated fraction of coinfections was reduced by approximately half, compared to the Palivizumab-free control (Fig. 5d). This suggests that there is a population of IAV that is dependent on RSV F to gain entry to cells, which in turn suggests that RSV F is the functional glycoprotein of IAV/RSV HVPs.

**Figure 5.**
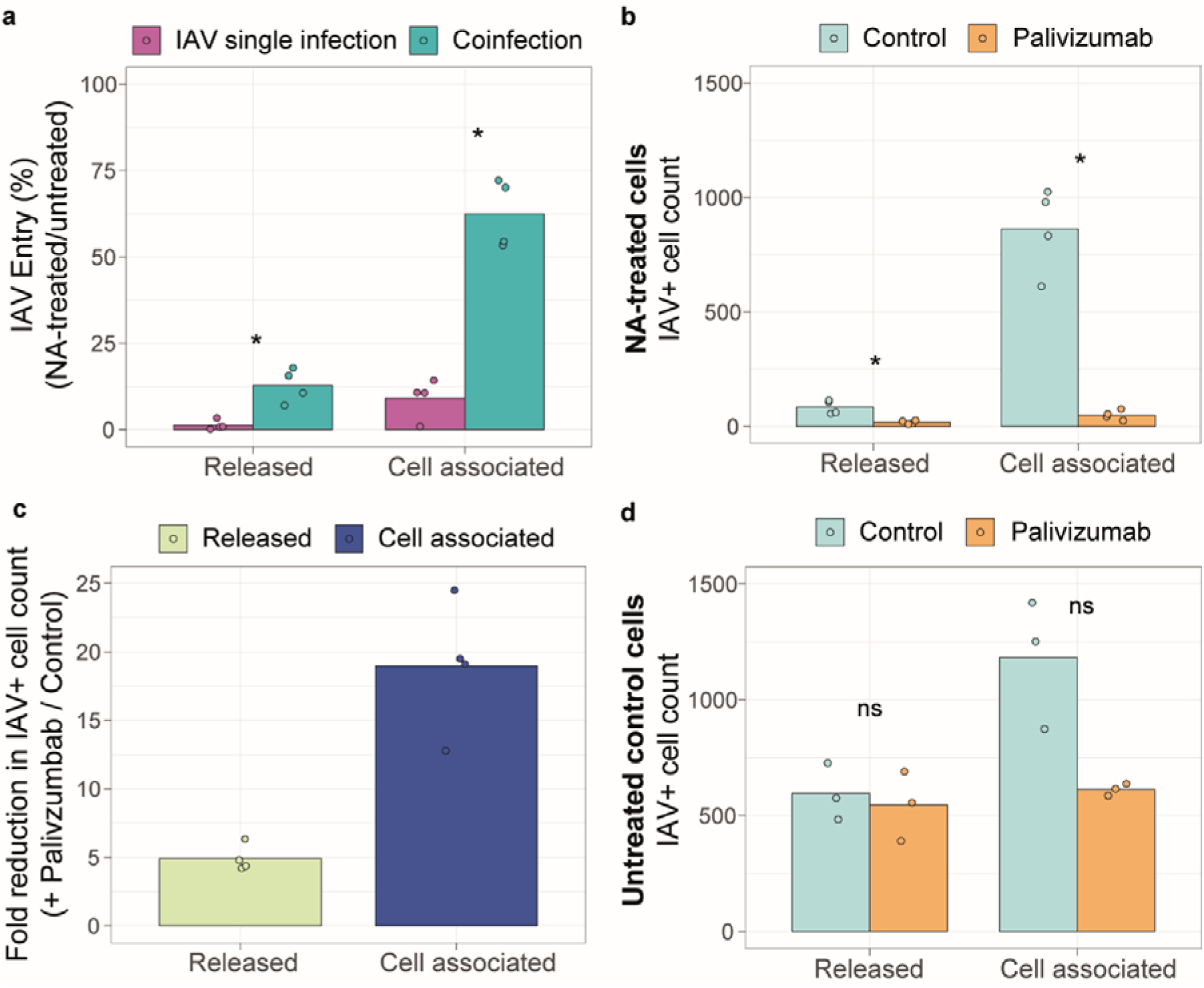
Virus produced from coinfection facilitates IAV infection in sialic acid-depleted cells. NA-treated or control A549 cells were infected with virus harvested from coinfection or IAV single infection, fixed and immunostained at 12 hpi and the number of IAV infected cells was quantified using an automated image-based cell counter. (a) Ratio of IAV entry into NA-treated cells versus control cells when harvested from single infection (magenta bars) or mixed infection (teal bars). IAV entry to NA-treated cells was calculated as a percentage of the IAV-positive cell count in the matched untreated control. (b) IAV positive cell count in NA-treated cells infected with virus that was harvested from coinfection and incubated with palivizumab (orange bars) or a serum free control (cyan bars). (c) Fold change in IAV positive cell counts as shown in (e) for released (yellow bar) and cell associated (navy bar) virus. (d) IAV positive cell count in untreated control cells infected with virus that was harvested from coinfection and incubated with palivizumab (orange bars) or a serum free control (cyan bars). Data was collected from three independent experiments and statistical significance was determined by Mann Whitney test, * indicates p<0.05 and ns indicates p>0.05.

Our cryo-ET data showed that HVPs contain IAV and RSV genomes. To determine if they possess infectivity for both viruses, we infected NA-treated cells with virus harvested from single or mixed infections. At 12 hpi, cells were fixed and stained for IAV HA and RSV F and imaged by confocal microscopy. The presence of coinfected cells suggested that both genomes were delivered into the cells simultaneously, likely by HVPs (Fig. 6a-b). This conclusion is based on the facts that 12 hours is not a sufficiently long time to allow extensive cell-to-cell spread by IAV and RSV which may result in coinfection; and because the probability of coinfection by chance was low as the MOI for each virus was approximately 0.01 (based on back-titrations of harvested virus). Examination of coinfected cells using super-resolution microscopy revealed viral IAV HA and RSV F double-positive filaments (Fig. 6b), suggesting that the HVPs can be maintained over virus passage. To establish if HVPs could spread IAV from cell to cell within a population of cells refractory to IAV infection, we infected NA-treated or control cells with virus harvested from IAV single infections or from the cell-associated fraction of mixed infections (the fraction with enriched HVPs). We then applied an overlay to prevent virion diffusion and incubated the cells for 48 hours. As expected, in single IAV infections, abundant IAV-positive cells were observed in non-treated cells (Fig. 6c, top row). In the NA-treated cells, no infection by IAV from single infections was observed (Fig. 6c, middle row), whereas IAV foci consisting of multiple distinct infected cells were observed in NA-treated cells infected with virus from mixed infections (Fig. 6c, bottom row). Notably, IAV-positive foci colocalised with RSV-positive foci (Fig. 6c, bottom row). These results suggests that HVPs can mediate the spread of IAV within a refractory cell population.

**Figure 6:**
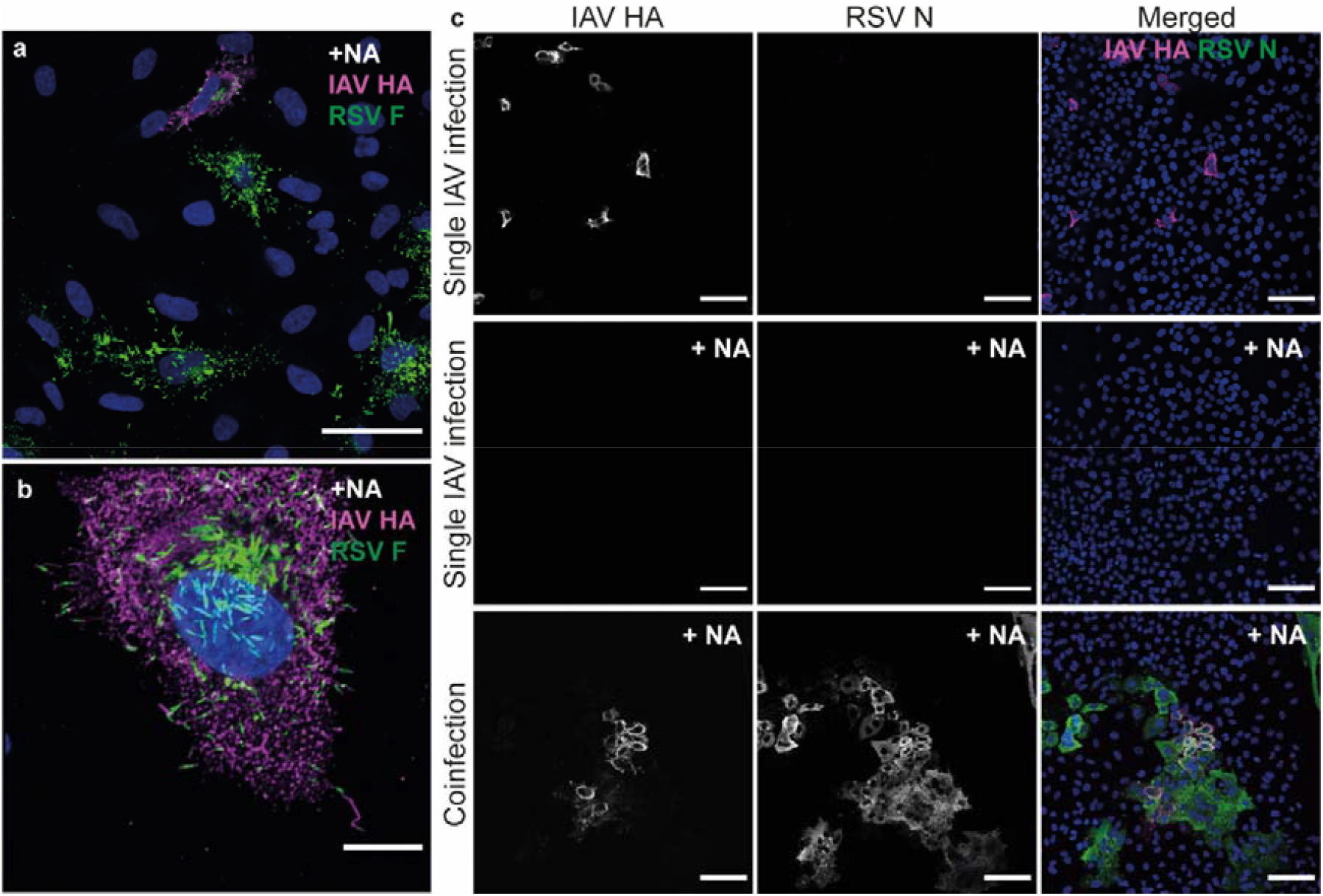
Hybrid viral particles possess dual infectivity for IAV and RSV and can facilitate spread of IAV in a refractory cell population. (a) A549 cells were infected with cell-associated virus from a coinfection and incubated for 12 hpi, before staining for IAV HA (magenta) and RSV F (green) and imaging by standard confocal microscopy. Scale bar indicates 100 µm. (b) Super-resolution confocal imaging of coinfected cells shows formation of dual-positive filamentous structures, indicating hybrid particle formation is maintained upon passage. Scale bar indicates 1 µm. (c) Virus harvested from IAV single infection (top and middle rows) or coinfection (bottom row) was incubated on untreated control wells (top row) or neuraminidase-treated wells (middle and bottom rows) with an overlay for 48 hours. Foci of infection were immunostained for IAV HA (magenta) and RSV N (green) and imaged by confocal microscopy. Infection was observed by IAV from single infection in untreated control cells, but not neuraminidase-treated cells. Virus harvested from coinfection formed large foci and IAV foci colocalised with RSV foci. Scale bar indicates 200 µm.

To determine the potential for HVP formation in a more biologically relevant system, we coinfected differentiated primary human bronchial epithelial cells (HBECs). We observed no difference in IAV replication kinetics between single-infected and coinfected cultures (Fig. 7a). In coinfections, RSV titres were variable, but the average titres over experimental replicates were lower than in single infections (Fig. 7b). Trends in replication kinetics between single and mixed infections matched the trends observed in A549 cells, providing confidence that viral interactions are conserved between the two systems. To examine the spread of IAV and RSV in HBECs, we fixed and paraffin-embedded infected or mock-infected cultures (culture morphologies shown in Fig. 7c), and performed immunostaining using antibodies targeting IAV HA, IAV NP and a polyclonal antibody raised against the whole RSV virion. We observed diffuse staining for IAV HA and RSV across the apical layer of cells in single infections, whereas in coinfected cultures there was a high degree of individual cells infected by each virus as well as evidence of coinfected cells (Fig. 7d). Analysis of individual coinfected cells revealed dual staining for both IAV and RSV antigens both at the apical surface of coinfected cells and within the cytoplasm (Fig. 7e-f), providing an opportunity for interactions between IAV and RSV and the formation of HVPs.

**Figure 7:**
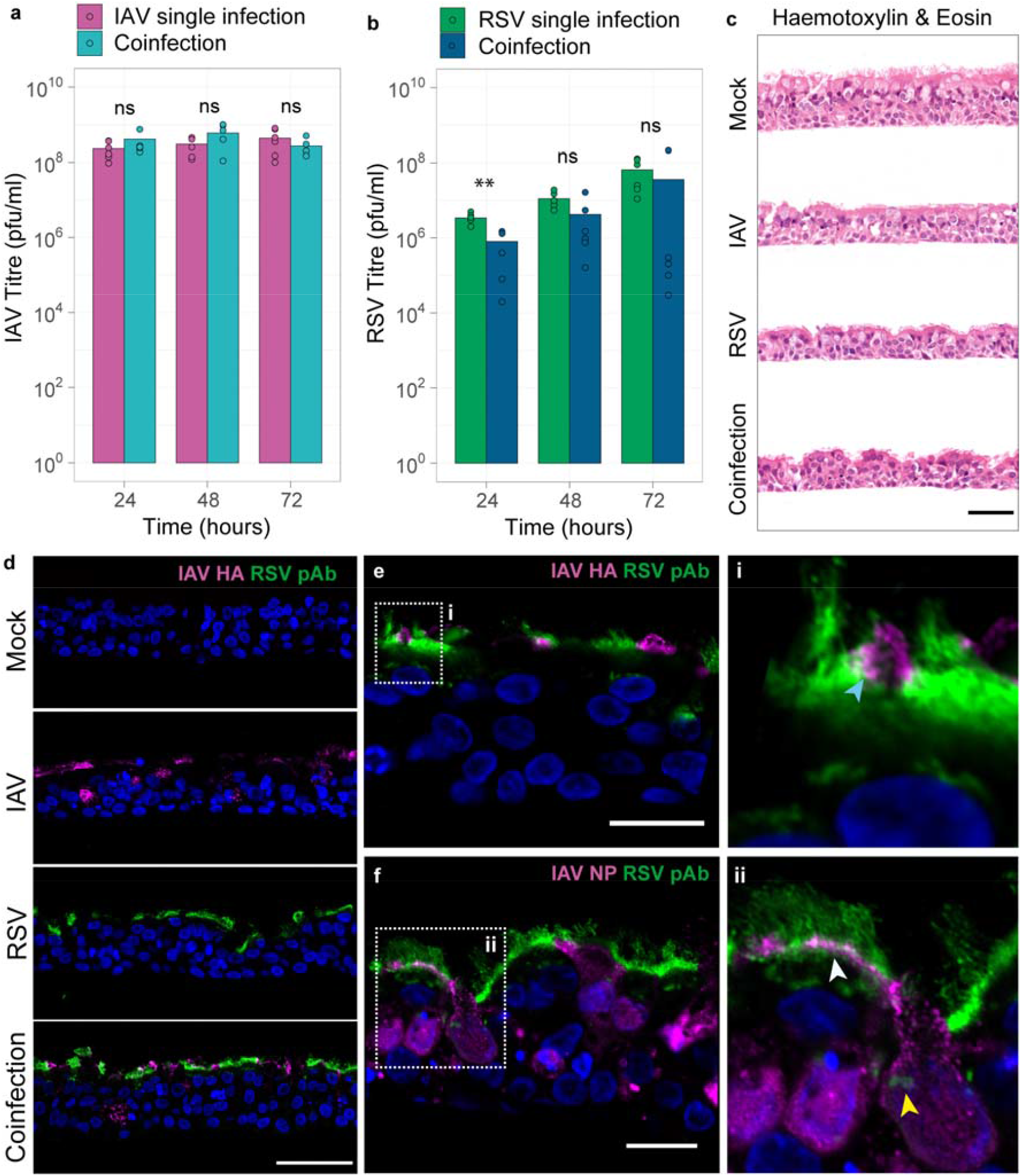
Coinfection of primary differentiated human bronchial epithelial cell cultures (hBECs) results in robust replication of IAV and RSV, with evidence of cellular coinfection. (a) IAV replication kinetics in hBEC cultures in single infection (magenta bars) and mixed infection with RSV (teal bars), n=6 individual transwells over 3 independent experiments. (b) RSV replication kinetics in hBEC cultures in single infection (green bars) and mixed infection with RSV (dark blue bars). n=6 individual transwells over three independent experiments. (c) Paraffin embedded sections collected at 24 hpi, stained with haemotoxylin and eosin. (d) Immunofluorescence showing IAV HA (magenta) and RSV antigen (green) localisation in paraffin embedded sections at 24 hpi. (e) Coinfected section imaged by super-resolution confocal microscopy shows localisation of IAV HA (magenta) and RSV antigen (green) to the apical surface of epithelial cells. Scale bar represents 20 µm. The magnified area (i) shows the region indicated by the white dashed box, highlighting a coinfected cell with IAV HA and RSV antigen localised at apical surface (blue arrow). (f) Localisation of IAV NP (magenta) and RSV antigen (green) within coinfected cells. Scale bar represents 20 µm. The magnified area (ii) shows the region indicated by the white dashed box highlighting IAV NP and RSV antigen within the cytoplasm, where RSV inclusion bodies can be observed (yellow arrow), and at the apical surface of coinfected cells (white arrow).

## Discussion

Respiratory viruses share a common tropism for the human respiratory tract and cause significant disease burden. While there is increasing evidence that interactions amongst viruses play an important role on virus dynamics and transmission, most of what is known about virus biology and pathogenesis is based on a tractable but reductionist research approach whereby each virus is studied in isolation.

Recent work provided evidence that interactions among respiratory viruses occur and have measurable outcomes at multiple levels, from populations, to individuals and tissues (*24-28*). However, studies characterizing direct virus-virus interactions within cells are scarce. Here we describe previously unknown interactions between IAV and RSV, two clinically important respiratory viruses that belong to different taxonomical families.

We show that, in coinfections IAV replicates to equivalent or marginally higher titres compared to single IAV infections, whereas RSV replication is reduced. The consistency in IAV replication kinetics in the presence or absence of RSV contrasts with the inhibition of IAV replication in coinfections with rhinovirus (*26*). This indicates that the consequences of coinfections are highly dependent on the viruses involved as they trigger virus-specific cellular responses.

We also show compelling evidence that coinfections can generate infectious HVPs composed of structural, genomic, and functional components of both parental viruses. As HVPs can evade IAV-targeted neutralisation and infect cells lacking IAV receptors suggests that coinfections can generate viruses with altered antigenicity and expanded tropism. Using Palivizumab, we showed that RSV F mediates HVP entry, indicating that in the context of a hybrid particle, IAV can use the glycoprotein of an unrelated virus as its functional envelope protein. This property may facilitate within-host dissemination to areas of the respiratory tract that are refractory to infection by one of the parental viruses, which is likely to impact pathogenesis and disease outcome. For example, IAV predominantly infects the upper and middle respiratory tract causing uncomplicated influenza, while RSV spreads more readily to the middle and lower respiratory tract (LRT) (*29, 30*). HVPs could enable IAV to escape mucosal antibodies while spreading to the LRT, with subsequent potential complications, including viral pneumonia (*30*). In addition, as IAVs exhibit high mutation rates, LRT infections by HVPs might favor the selection of IAVs with increased tropism for the LRT and therefore result in selection of more pathogenic viruses.

In recent years, a novel conceptual framework that incorporates social evolution theory has been developed to explain how virus-virus interactions can play an important role on virus function and fitness (*31*). We show that HVP formation is maintained over multiple rounds of infection and that HVPs facilitate the spread of IAV within a population of refractory cells. This observation aligns with the concept that, like other pathogens and organisms, viruses can engage in social-like traits, that are beneficial to virus fitness and function.

Using a lung-derived human cell line, we show that the generation of HVPs by coinfection is biologically feasible. The fact that IAV and RSV cocirculate in winter in the same populations (*1*), have a shared tropism for ciliated epithelial cells (*32, 33*), bud from the apical cellular surface (*22, 33, 34*) and coinfect cells within the respiratory epithelium (this work) suggest that HVPs have the potential to be generated *in vivo*. The likelihood of a cell becoming coinfected during natural infection remains unknown but will vary depending on the timing of infection and the localization of infectious foci within the respiratory tract. Estimates of viral bursts show that as viral load increases, the effective MOI to susceptible cells surrounding an infectious focus increases (*35*), enhancing the probability of cellular coinfection and therefore the potential generation of HVPs.

The formation of HVPs raises questions about fundamental rules that govern viral assembly and budding. These processes, which are thought to be highly regulated, involve selective recruitment, trafficking (*36, 37*) and multimerization of viral proteins (*19, 38-40*) within specific compartments of the cell. While we described the formation of HVPs as a consequence of coinfection by IAV and RSV, we hypothesize that coinfections involving other pleomorphic enveloped viruses are also likely to generate HVPs. However, we pose that formation of infectious HVPs requires more than structural compatibility, and includes similar tropism, absence of superinfection exclusion or interference, as well as seasonal and geographical co-circulation. RSV is a pleomorphic enveloped virus with a broad tropism for different regions of the respiratory tract and is frequently observed in co-infections in winter (*41-43*), therefore RSV is a good candidate to form HVPs with other respiratory viruses. This might explain some of the mechanisms that lead to viral pneumonia (*44*). Further studies will be required to address which virus combinations can generate infectious HVPs; which viral properties favor their formation; and how they impact on pathogenesis and virus transmission.

## Online Methods

### Cells

Human lung adenocarcinoma cells (A549) (American Type Culture Collection [ATCC], CCL-185), Madin Darby canine kidney (MDCK) cells (ATCC, CCL-34) and HEp-2 (ATCC, CCL-23) cells were cultured in Dulbecco’s Minimum Essential Media (DMEM), high glucose, GlutaMAX supplemented with 10% fetal bovine serum (FBS).

Human bronchial epithelial cells (hBEC) were cultured and differentiated at air-liquid interface as describe previously (*24*). Briefly, human bronchial epithelial cells (hBEC) (Epithelix,) were cultured in Epithelix human airway epithelial cell medium (Epithelix; EP09AM) 37°c, 5% CO2 in a humidified incubator. Cells were cultured in tissue culture flasks until 80% confluent. After this point, cells were trypsinised and seeded at 2×10^4^ cells/transwell onto transwell inserts for 24-well plate with 0.4 µm pore size with a pore density of 1.6×10^6^ pores per cm2 (Falcon, 734-0036). When cells were fully confluent on transwell membranes, apical media was removed to initiated air-liquid interface (ALI). Basal media was replaced with Pneumacult-ALI media (STEMCELL Technologies, 05001). Basal media was replenished every 2-3 days. When cultures began producing mucus (approximately 20 days post ALI initiation), the apical surface of cultures was washed twice weekly with serum free DMEM.

### Viruses

H1N1 influenza A/Puerto Rico/8/34 was rescued as previously described (*45*) and virus stocks were grown in MDCK cells. RSV strain A2 (American Type Culture Collection, VR-1540) was grown in HEp-2 cells.

### Antibodies

The following primary antibodies were used at optimised concentrations: RSV fusion glycoprotein (Abcam, UK, AB24011, 1/1000), RSV nucleoprotein (Abcam, UK, AB22501, 1/1500), antisera to influenza A H1 (A/Puerto Rico/8/34, 1/1000) (NIBSC, UK, 03/242), influenza A virus nucleoprotein (European Veterinary Society, EVS238, 1/1000), mouse monoclonal anti-IAV (A/Puerto Rico/8/34) HA (Sinobiological, 11684-MM03, 1/500), polyclonal anti-IAV (A/Puerto Rico/8/34) nucleoprotein kindly provided by Paul Digard, goat polyclonal anti-RSV (Abcam, AB20745, 1/500) Secondary antibodies: Rabbit anti-mouse IgG alexafluor 488 conjugate (Sigma-Aldrich, USA, SAB4600056, 1/1000), donkey anti-mouse 594 conjugate (Abcam, UK, ab150108, 1/1000) donkey anti-sheep alexafluor 594 conjugate (Thermo Fisher Scientific, USA, A-11016, 1/1000),), goat anti-rabbit alexafluor 488 (Invitrogen, A11034).

### Virus growth curves

A549 cells were infected with IAV, RSV or synchronously infected with a mixed inoculum of IAV and RSV, diluted in DMEM, with 2% FBS and 1µg/ml trypsin TPCK. Infections were carried out with RSV at a multiplicity of infection (MOI) of 4 and IAV at an MOI of 4 or 0.4. A549 cells were inoculated in 48-well plates and incubated at 37°C, 5% CO2 for 90 minutes, before the inoculum was removed and replaced with DMEM, with 2% FBS and 1µg/ml trypsin TPCK. Cells were incubated at 37°C, 5% CO2 and supernatant from each infection was collected at 24, 48 and 72 hours post infection and stored at −80°C prior to titration by plaque assay.

HBEC cultures were infected no earlier than 35 days post ALI initiation. The apical surface of cultures was washed with DMEM 24 hours prior to infection by applying pre-warmed DMEM to the apical surface of cultures and incubating at 37°c, 5% CO2 in a humidified incubator for 20 minutes, followed by removal. This washing step was repeated immediately before infection. Inoculum containing 105 pfu of IAV, RSV or a mixed inoculum of both viruses (105 pfu of each virus) was prepared in DMEM. Cultures were incubated with inoculum for 2 hours at 37°c 5% CO2, after which the inoculum was removed and cultures were washed once with DMEM as described. Inoculum was back titrated back plaque assay to confirm virus input and served as the zero hours time point for growth curves. Two cultures were infected per infection condition and experimental time point. Samples were collected from the apical surface of cultures at 24, 48 and 72 hpi, by incubating with DMEM for 30 minutes. Sample were then removed and stored at −80°c, prior to titration by IAV or RSV plaque assay. Transwells were fixed with 4% formaldehyde for 1 hour. After fixation, HBEC cultures were embedded in paraffin blocks, cut to 2-3 µm thick sections and mounted on glass slides. Sections were stained with haematoxylin and eosin to determine morphology.

Infectious titre was determined by plaque assay in MDCK cells or HEp-2 cells for IAV and RSV, respectively. Validation was carried out prior to these experiments to ensure specific detection of IAV or RSV plaques in the respective cell type. While IAV and RSV are capable of infecting both cell lines, each cell line is only permissive to plaque formation by one virus I.e. IAV forms plaques in MDCK but not HEp-2cells, while RSV forms plaques in HEp-2 but not MDCK cells. Viruses were titrated in 10-fold serial dilutions in duplicate and incubated on confluent cell monolayers for 90 minutes. For IAV plaque assays, overlay containing DMEM, 1µg/ml trypsin TPCK and 0.6% Avicel was used and plates were incubated at 37°C, 5% CO2 for 48 hours. For RSV plaque assays, overlay containing DMEM, 5% FBS and 0.6% Avicel was applied and plates were incubated at 37°C, 5% CO2 for 4-5 days. Plates were then fixed with 4% formaldehyde in phosphate buffered solution (PBS) and stained with 0.1% Coomassie Brilliant Blue. Plaques were counted and titre determined as plaque forming units per ml (pfu/ml). Experiments were carried out in technical (n=3) and biological triplicate (n=3 independent experiments).

### Immunofluorescence and confocal microscopy

Cells were seeded at 2×10^5^ cells/ml on 13 mm glass coverslips or 35mm glass bottom dishes (MATTEK Corporation Inc, USA) prior to infection. Infections were incubated for 24 hours prior to fixation with 4% fomaldehyde in PBS. Fixed cells were permeabilized with 0.1% triton X100 for 10 minutes at room temperature. Samples were blocked with 1% bovine serum albumin (BSA) in PBS for 30 minutes. Primary and secondary antibodies were diluted in 1% BSA in PBS and incubated for 60 minutes at room temperature, followed by four washes with PBS. Coverslips were mounted with Prolong Gold mounting media with DAPI (Invitrogen, USA, P36392), and samples in dishes were stained with 2 µg/ml Hoescht 33342 solution for 10 minutes. For combining mouse primary antibodies, samples were blocked in rabbit serum (Gentex, USA, GTX73221), then stained with primary antibody and rabbit anti-mouse secondary antibody, followed by secondary blocking in donkey serum (Gentex, USA, GTX73205), then stained with primary antibody and donkey anti-mouse secondary antibody. For live-cell imaging, cells were incubated with 2 µg/ml Hoescht 33342 solution in cell culture media, for 30 minutes at 37°c. Cells were then washed with cold PBS, followed by incubation with primary antibody diluted in 2% BSA in PBS for 5 minutes on ice. Cells were washed three times with cold PBS and incubated with secondary antibody diluted in 2% BSA in PBS for 5 minutes on ice. Cells were stored in PBS and imaged immediately following staining.

For immunofluorescence staining, sections were dewaxed by heating in an oven at 60°c for 1 hour. Next, slides were washed four times with xylene, each for 10 minutes. Following this, sections were rehydrated via washes with 1:1 (v/v) Xylene:Isopropanol mixture, then 100%, 90%, 70% and 50% isopropyl alcohol solution each for five minutes. Sections were washed thoroughly with d.H_2_O and PBS. For antigen retrieval, sections were treated with proteinase K solution (Dako, S3020) for 15 minutes.

Sections were mounted into humid chambers for immunostaining. Sections were permeablised with 1% triton X100 for 10 minutes, followed by three washes with PBS. Sections were blocked with 2% BSA in PBS for 30 minutes at room temperature. Primary antibodies were diluted in 2% BSA solution and applied to sections for two hours at room temperature, followed by washing with PBS. Secondary antibodies were diluted in 2% BSA solution and applied to sections for 1 hour in the dark at room temperature, followed washing with PBS and d.H_2_O. Slides were mounted with Prolong Gold mounting media containing DAPI (Invitrogen, P36392).

Confocal microscopy was carried out using Zeiss LSM880 AxioObserver microscope (ZEISS, Germany). Standard images were collected using GaAsP detector with 405nm, 488nm and 598nm excitation lasers, using 40x/1.4 or 63x/1.4 plan-apochromat oil DIC M27 objectives. Super-resolution imaging was collected using the Airyscan detector with 405nm, 488nm and 594nm excitation lasers, using plan-apochromat 63x/1.4 oil DIC M27 objective. Z-stacks were collected using a z-interval of 100nm. For determining proportions of infection and focus assay on neuraminidase treated cells, fields of view were imaged using 3×3 tile scans at 40x/1.4 oil DIC M27 objective. Image processing and analysis was carried out in ImageJ software (v2.0.0-rc-56/1.52g) (NIH, USA) and Zeiss Zen lite (Blue edition) version 2.6. Single or coinfected cells were counted manually, using cell counter plugin on ImageJ. Experiments were carried out in technical (n=3) and biological triplicate (n=3 independent experiments).

### Scanning electron microscopy

Cells were seeded at 2×10^5^ cells/ml on 13 mm glass coverslips prior to infection at MOI of 4 with IAV, RSV or coinfection for 24 hours. Cells were fixed with 1.5% Glutaraldehyde in 0.1M Sodium Cacodylate (SC) buffer for 1 hour and washed 3 times for 5 minutes each with 0.1M SC buffer before incubation with 1% osmium tetroxide for 1 hour. Samples were then stained with aqueous 0.5% Uranyl Acetate for 1 h and further dehydrated through an ethanol series and subjected to critical point drying using an Autosamdri-815 Critical Point Dryer (Tousimis, USA) before mounting and coating in 20nm Gold/Palladium using a Quorum Q150T ES High vacuum coating system (Quorum Technologies, UK). Images were collected using JEOL IT100 SEM at 20kV, with InTouch Scope software, version 1.05 (JEOL USA Inc., USA). Infections and imaging were carried out in two independent experiments, with duplicate samples per experiment.

### Cryo-electron Tomography and computational analysis

Cells were seeded at 2×10^4^ cells/ml on laminin-coated (Roche) 200-mesh gold R2/2 London finder Quantifoil grids (Electron Microscopy Sciences) in 35 mm glass bottom dishes (MATTEK Corporation Inc, USA). Cells were infected with IAV MOI 1 and RSV MOI 4 and incubated for 24 hours at 37°C. Cryo-ET imaging experiments of coinfected cells were carried out in three independent experiments, with duplicate grids imaged per experiment. Immediately before plunge-freezing 3 µl colloidal solution of 20 nm gold fiducials (Sigma-Aldrich) pretreated with BSA was added to each grid. The gold served as fiducial markers for cryo-tomogram reconstruction. Grids were blotted from the backside of the grid and plunge-frozen into liquid ethane using the FEI Vitrobot Mark IV (Thermo Fisher Scientific, USA) or Leica EM GP 2 (Leica Microsystems, Germany). Plunge-frozen grids were subsequently loaded into a JEOL CRYO ARM 300 (JEOL Ltd, Japan) equipped with an energy-filter and DE64 detector (Direct Electron, USA). To identify the presence of virus budding sites near electron transparent cellular edges low-magnification grid maps were generated using serialEM (*46*). Next, polygon maps at 3,000x magnification were collected at areas of interest and virus budding sites were identified. Tilt-series of virus budding sites were then recorded using SerialEM at either 30,000x (1.921 Å/pixel) or 50,000x (1.499 Å/pixel) magnification. Each tilt-series was collected from −60° to +60° at an increment of 2° or 3° at 5-to 8-µm underfocus (30,000x) and 2-to 5-µm underfocus (50,000x). The cumulative dose of one tilt-series was between 80 and 120 e^−^/Å^2^. Tilt-series were collected using a dose-symmetric scheme starting at 0° implemented in SerialEM (*47*) and were collected in ‘movie mode’. For the 30,000x magnification tilt-series movie frames were aligned using alignframes module in IMOD (*46*). For the 50,000x magnification tilt-series movie frames were aligned using MotionCor2 with 6 by 6 patches (*48*). Once tilt-series sub-frames were aligned and binned eightfold into 1 k × 1 k arrays they were reconstructed into 3D tomograms using weighted back projection with IMOD. To aid interpretation and visualization, noise reduction was applied using Topaz (*49*). Figures and supplemental videos were prepared using IMOD, Illustrator ImageJ and Quicktime. Segmentation and isosurface rendering were performed manually using the thresholding tool in Amira (Thermo Fisher Scientific, USA).

Glycoprotein-distance was measured and calculated using the imodinfo module in IMOD. Denoised tomograms were averaged between 5-10 slices to improve contrast and glycoproteins were viewed from top down. Distances were measured between the centres of pairs of adjacent glycoproteins. For IAV and RSV controls, spike measurements were collected from 11 individual tomograms for each virus (n=326 for IAV, n=236 for RSV). For pseudotyped virus, spike measurements were collected from each particle, from 4 individual tomograms (n=50 measurements per tomogram).

### Neutralisation assays

A549 cells were seeded at 1×10^5^ cells/well in 24-well plates prior to infection with IAV (MOI of 1), RSV (MOI of 4) or mixed infection. Infected cells were incubated for 24 hours before released virus was harvested from supernatant and cell associated virus was collected by scraping and vortexing infected cells. Virus stocks were back titrated in MDCK or HEp-2 cells to determine pfu/well in the neutralisation assays. For IAV neutralisation assay, IAV from single infection or mixed infection was incubated with polyclonal anti-serum targeting A/Puerto Rico/8/34 H1 (NIBSC, UK, 03/242) at 1/200 dilution in DMEM or a serum-free control. For the RSV neutralisation assay, RSV from single infection or mixed infection was incubated with Palivizumab (Evidentic, Germany) at 10 µg/well concentration diluted in DMEM, or a serum-free control. Virus (neat or at a 1/10 dilution in DMEM) was incubated with serum for 1 hour at 37°c before transfer to A549 cells that had been seeded at 1×10^4^ cells/well in 96-well plates 48 hours prior to infection. Infections were incubated for 12 hours at 37°c and following this, cells were fixed in 4% fomaldehyde. Plates were immunofluorescence stained for IAV or RSV nucleoprotein and infected cells were quantified using the Celigo automated cytometer (Nexcelom Bioscience, USA), using two target expression analysis. Neutralisation was calculated from the positive cell count in each technical replicate of the serum containing wells as a percentage of the average positive cell count in the serum-free control. Experiments were carried out in biological triplicate (n=3 independent experiments).

### Neuraminidase assay

To prepare fresh virus stocks for the neuraminidase assay, A549 cells were seeded at 1×105 cells/well in 24-well plates prior to infection with IAV (MOI of 1), RSV (MOI of 4) or mixed infection. Infected cells were incubated for 24 hours before released virus was harvested from supernatant and cell associated virus was collected by scraping and vortexing infected cells. Virus stocks were back titrated in MDCK or HEp-2 cells to determine pfu/well in the neutralisation assays. For the neuraminidase assay, cells were seeded at 1×10^4^ cells/well in 96-well plates, prior to treatment with 1mU/µl neuraminidase from clostridium perfringens (Sigma-Aldrich, USA, N2876) for 2 hours. To confirm removal of sialic acids, neuraminidase treated and control cells were stained with biotinylated Maackia Amurensis Lectin II (MAL II) (Vector Laboratories, UK, B-1265-1), followed by fluorescein conjugated streptavidin (Vector Laboratories, UK, SA-5001-1) or Erythrina Cristagalli Lectin conjugated to fluorescein (Vector Laboratories, UK, FL-1141-5). Neat virus stocks were transferred directly onto neuraminidase treated or untreated cells in 96-well plates or on 13 mm glass coverslips and incubated for 12 hours. Virus harvested from coinfection was also incubated with 10 µg/well Palivizumab for 1 hour at 37°c, before transfer to NA-treated or untreated control wells and incubated for 12 hours. Cells were fixed and 96-well plates stained for IAV or RSV nucleoprotein, followed by rabbit anti-mouse 488 (Sigma Aldrich, USA, SABA4600056) secondary antibody. Infected cells were counted using Celigo automated cytometer (Nexcelom Bioscience, USA), using two target expression analysis. Viral entry ratio was calculated from the positive cell count in each technical replicate of the NA-treated wells as a percentage of the average positive cell count in the untreated control wells. Fixed coverslips were immunostained for IAV HA or RSV F and imaged by confocal microscopy using Zeiss LSM880 with or without Airyscan detector. Experiments were carried out in biological triplicate (n=3 independent experiments).

For the focus assay, cells were seeded onto 13 mm glass coverslips at 1×10^5^ cells/well. Cells were treated with 1mU/µl neuraminidase from for 2 hours, before infection with fresh virus stocks harvested from IAV single infection or coinfection or infection of untreated control cells. Infections were overlaid with DMEM containing 0.6% Avicel, 2% FBS and 1 µg/ml trypsin TPCK and incubated at 37°c for 48 hours. Following this, coverslips were fixed and immunostained for RSV N and IAV HA. Samples were imaged by confocal microscopy using Zeiss LSM880.

### Statistical analysis and Data Visualization

Statistical tests were carried out using Graphpad Prism, version 9.1.0. No assumptions about data normality were made and Mann Whitney test was used to determine statistical significance in growth kinetic experiments, neutralisation assays and neuraminidase assays. For inter-spike measurements on tomograms, normal distribution was tested using Shapiro-Wilk test and statistical difference between groups was determined by unpaired t-test. Data was visualised with RStudio version 1.3.1056 (*50*) using GGPlot2 package (*51*). Statistical significance was indicated in figures as ns p>0.05, * p<0.05, ** p<0.01, *** p<0.001, **** p<0.0001.

## Data Availability

Representative tomograms of the chimeric virus particles described in this paper have been deposited in the Electron Microscopy Data Bank (www.ebi.ac.uk/emdb) under accession codes EMD-13228 and EMD-13229.

## Author contributions

Author contributions are based on the CRediT taxonomy (https://casrai.org/credit/). J.H.: Investigation, Methodology, Formal analysis, Visualisation, Writing - original draft; S.V.: Investigation, Data curation, Formal analysis, Resources, Metodology, Validation, Visualisation, Supervision Writing-original draft; J.S.: Investigation, Writing – review and editing; K.D.: Investigation, Writing – review and editing; D.M.G.: Investigation, Writing – review and editing; M.C.: Investigation, Writing – review and editing; M.M.: Investigation, Methodology, Writing – review and editing; S.D.C: Formal analysis, writing, review and editing; D.B.: Resources, Funding acquisition, Writing – review and editing; P.R.M.: Conceptualization, Methodology, Validation, Data curation, Supervision, Funding acquisition, Project administration, Writing – original draft.

## Acknowledgments

This work was supported by grants from the Medical Research Council of the United Kingdom (MC_UU_12014/9 to PRM, MR/N013166/1 to JH, MR/R502327/1 to DMG, MC_UU_12014/7 to SV, and MC_UU_12014/7 to DB). The Scottish Centre for Macromolecular Imaging (SCMI) is funded by the MRC (MC_PC_17135) and SFC (H17007).

## Conflicts of interest

The authors declare no conflicts of interest.

Correspondence and requests for materials should be addressed to Pablo Murcia.

Reprints and permissions information is available at www.nature.com/reprints.

## Figures

**Movie 1**: Video showing serial sections through the z-axis of a tomogram of a hybrid particle with a multiple IAV region, formed during coinfection of IAV and RSV (corresponding image shown in Fig. 3). Glycoproteins and RNPs of both IAV and RSV are labeled and denoted by arrows.

**Movie 2**: Video showing serial sections through the z-axis of a tomogram of hybrid particle with IAV and RSV regions with an adjoining region with a clear lumen (corresponding image shown in Extended data Fig. 4C). Glycoproteins and RNPs of both IAV and RSV are labeled and denoted by arrows.

**Movie 3**: Video showing serial sections through a tomogram of a pseudotyped viral filament generated during IAV and RSV coinfection (corresponding image shown in Extended data Fig. 3). Glycoproteins and RNPs are labeled and denoted by arrows.

## Extended Data

**Extended Data Figure 1:**
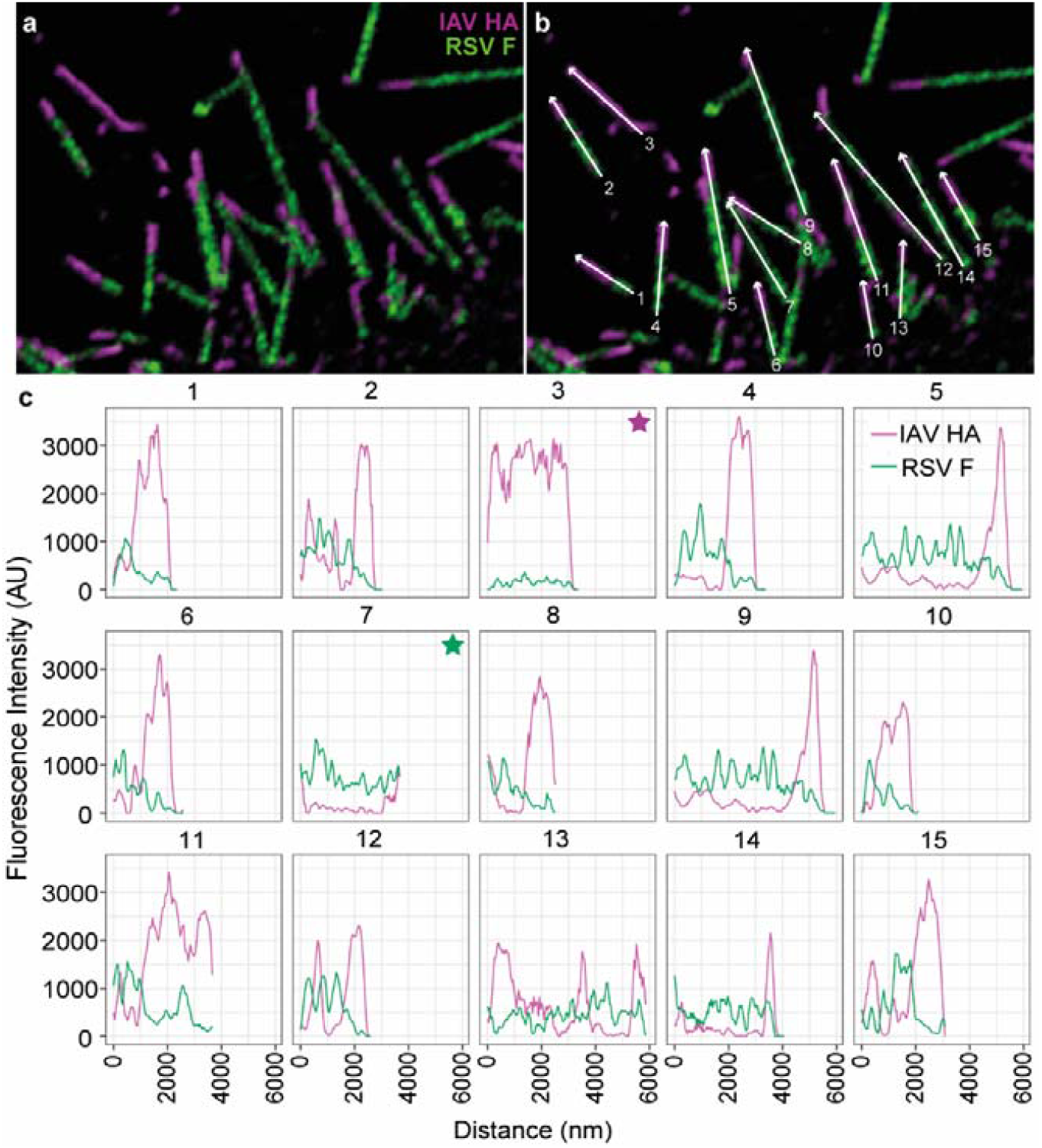
HA and F are expressed in discrete patches along the length of filaments, with HA predominantly at the distal end. (a) Magnified view of cell associated filaments (full image shown in figure 2A) show filaments with distinct patches of IAV HA (magenta) and RSV F (green) glycoproteins along the length of the filaments. (b) White arrows and filament numbering correspond to fluorescence intensity profiles displayed in (c). Minimal colocalisation was observed in the fluorescence intensity profiles (c) for IAV HA (magenta line) and RSV F (green line) signal along filaments numbered 1-15. IAV (filament 3, magenta star) and RSV (filament 7, green star) filaments were also identified among dual positive filaments.

**Extended Data Figure 2:**
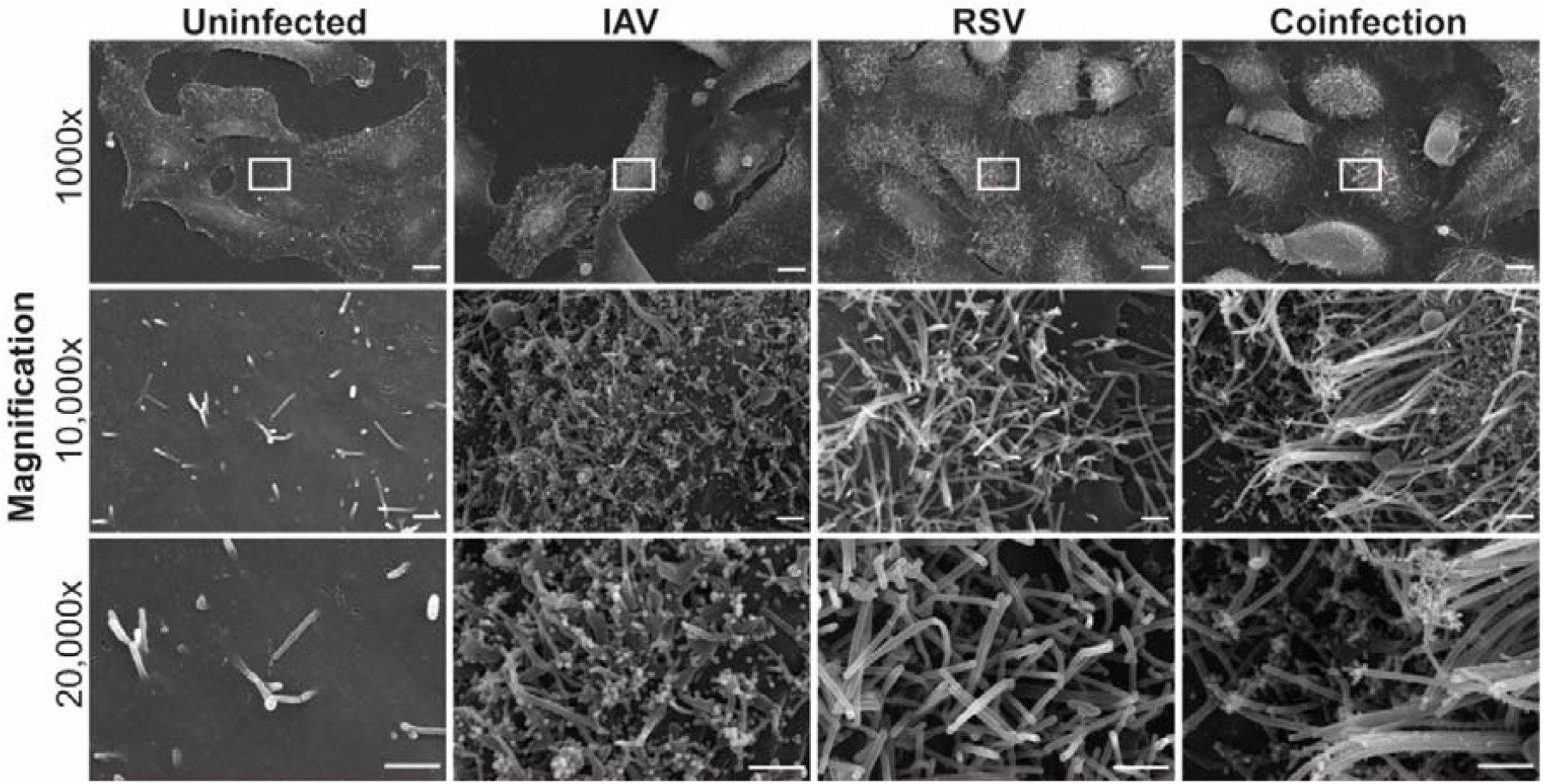
Scanning electron microscopy shows clear differences between IAV and RSV virion structure. Scanning electron micrographs of IAV, RSV, coinfected or mock infected cells imaged at 1000x (top row), 10,000x (middle row) and 20,000x (bottom row), region of magnification is denoted by the white box. Scale bars represent 10 µm at 1000x and 1 µm at 10,000x and 20,000x magnification.

**Extended Data Figure 3:**
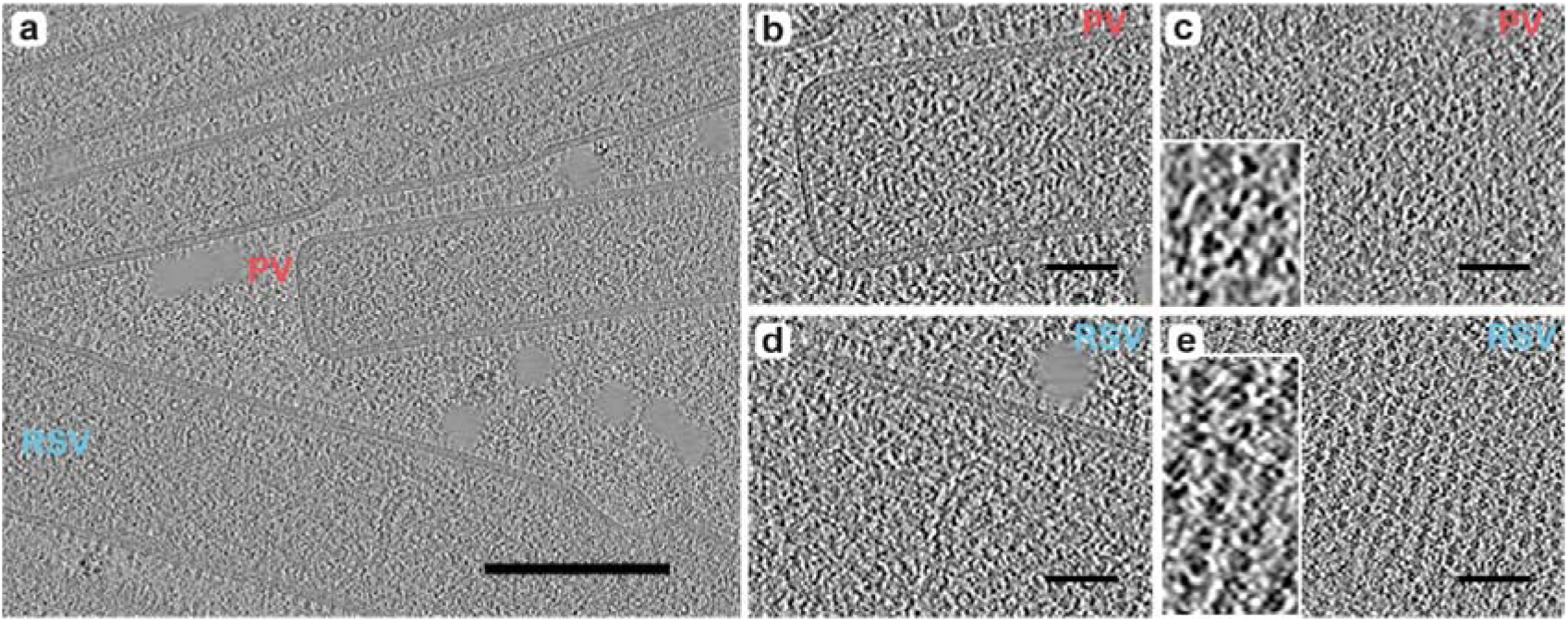
Coinfection generates RSV filaments that are pseudotyped with IAV envelope proteins. (a) Tomogram shows a pseudotyped RSV filament, indicated by red ‘PV’ label, near to RSV filaments, one example indicated by blue ‘RSV’ label. Scale bar indicates 200 nm. (b) Magnified cross-section of end of pseudotyped filament, showing RSV RNP contained within virion. (c) Surface of psuedotyped filament shows irregular arrangement of glycoproteins, with many displaying characteristic triangular shape of HA trimers, shown in magnified inset image. (d) Magnified cross-section of end of RSV filament, showing RSV RNP contained within virion and ultra-structure consistent with the pseudotyped virion. (e) Surface of the RSV filament shows helical arrangement of glycoproteins, with ring-shaped density of glycoproteins highlighted in magnified insert. Scale bars in panels (b-e) indicate 50 nm.

**Extended Data Figure 4:**
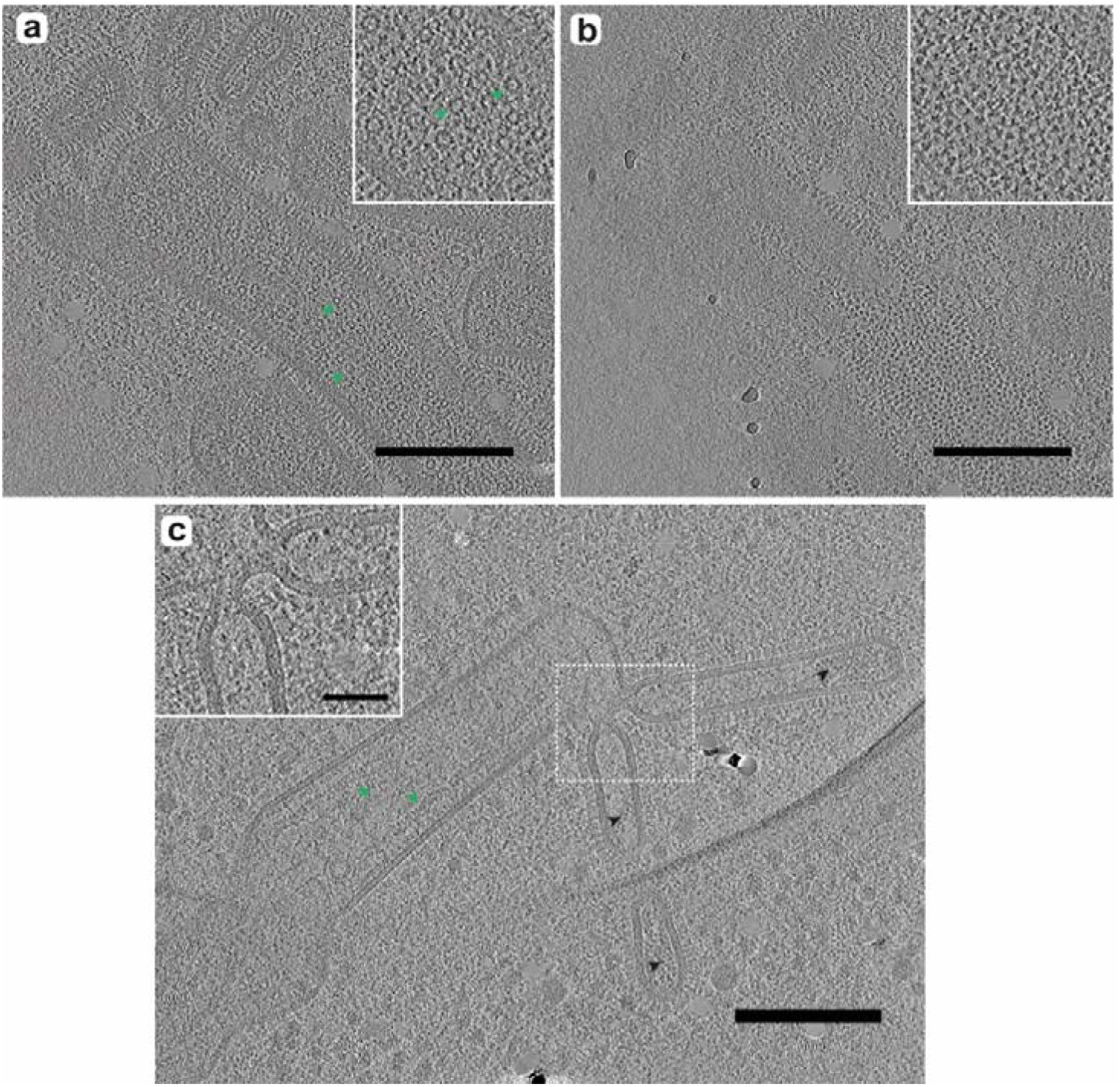
Further examples of hybrid particles. (a and b) show two z-positions through the same hybrid particle, which also displays pseudotyping in RSV-like region. (a) IAV-like regions extend from the top of the filament and ring-shaped densities corresponding to RSV genome, indicated by green arrows and highlighted in magnified inset image, are present within the virion. (b) The surface of the virion is covered in glycoproteins that are consistent in shape and arrangement with IAV glycoproteins, highlighted in magnified inset. Scale bars indicate 200 nm. (c) Tomogram shows a further example of a hybrid particle with two IAV-like regions which are joined to the RSV-like region by a continuous membrane. Black and green arrows indicate IAV and RSV RNP respectively, contained with in their associated structural regions. Scale bar indicates 200 nm. There is a clear shared lumen which continues between RSV and IAV regions, highlighted within magnified inset which corresponds to region marked by white dashed box. Scale bar indicates 50 nm.

**Extended Data Figure 5.**
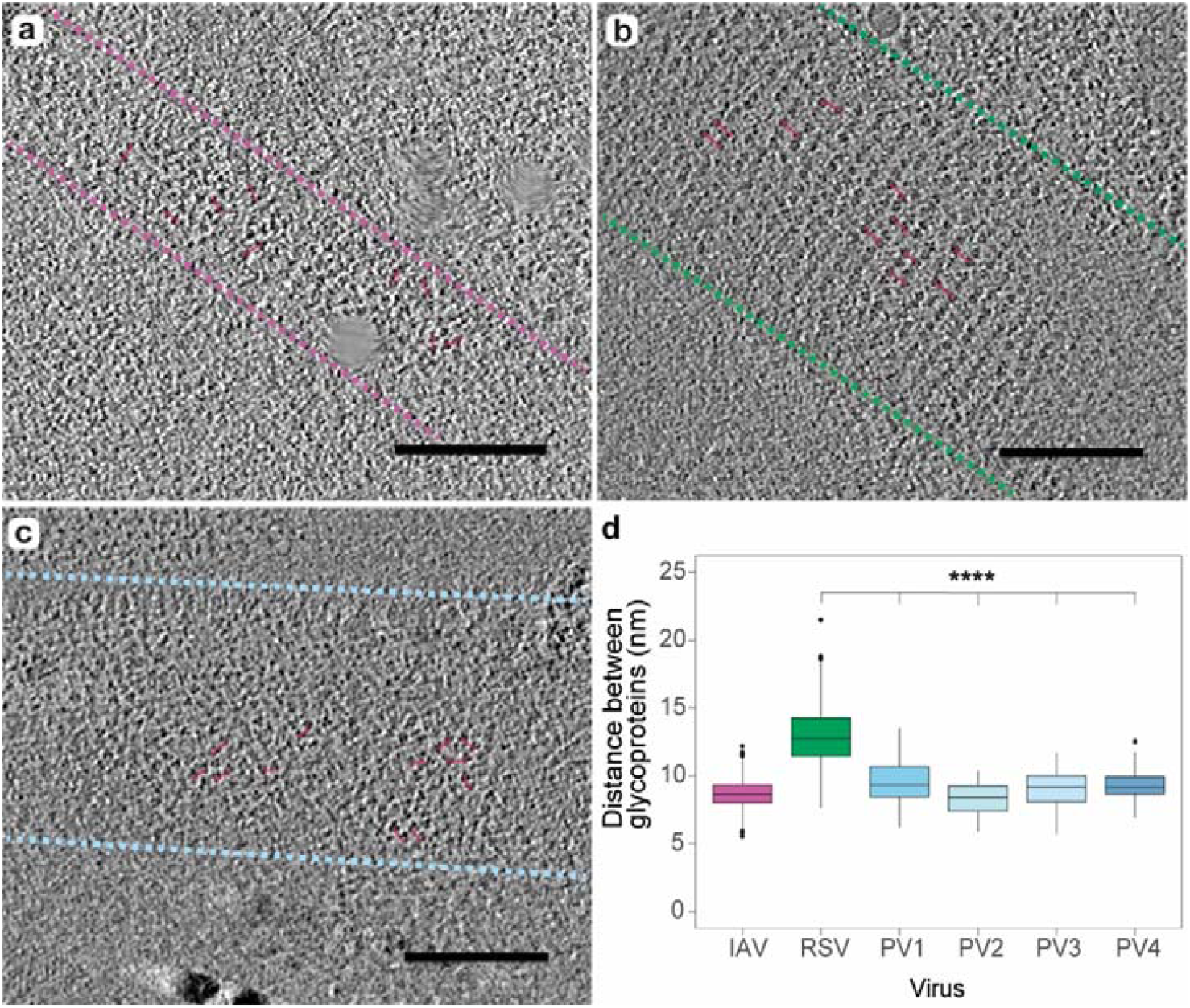
Inter-spike distance measurements reveal that pseudotyped viruses are decorated with IAV glycoproteins. To determine the glycoprotein arrangement on pseudotyped viruses, inter-spike distances were measured between glycoprotein pairs. Representative examples are shown for IAV (a), RSV (b) and pseudotyped virions (c) with red lines indicating example distances measured. Pink, green and blue dashed lines indicate the edges of IAV, RSV and psuedotyped filaments respectively. Scale bars indicate 200 nm. Control measurements were collected from 11 tomograms for IAV (measurements n=326) and 11 tomograms for RSV (measurements n=236). Measurements of pseudotyped virions were collected from 4 individual tomograms (n=50 measurements per tomogram). (d) Unpaired t-test analysis confirmed glycoproteins distance measurements were significant different between RSV and pseudotyped virions, with average inter-spike distances of 8.71 nm for IAV, 12.9 nm for RSV and a range of 8.31-9.56 nm for pseudotypes.

**Extended data figure 6:**
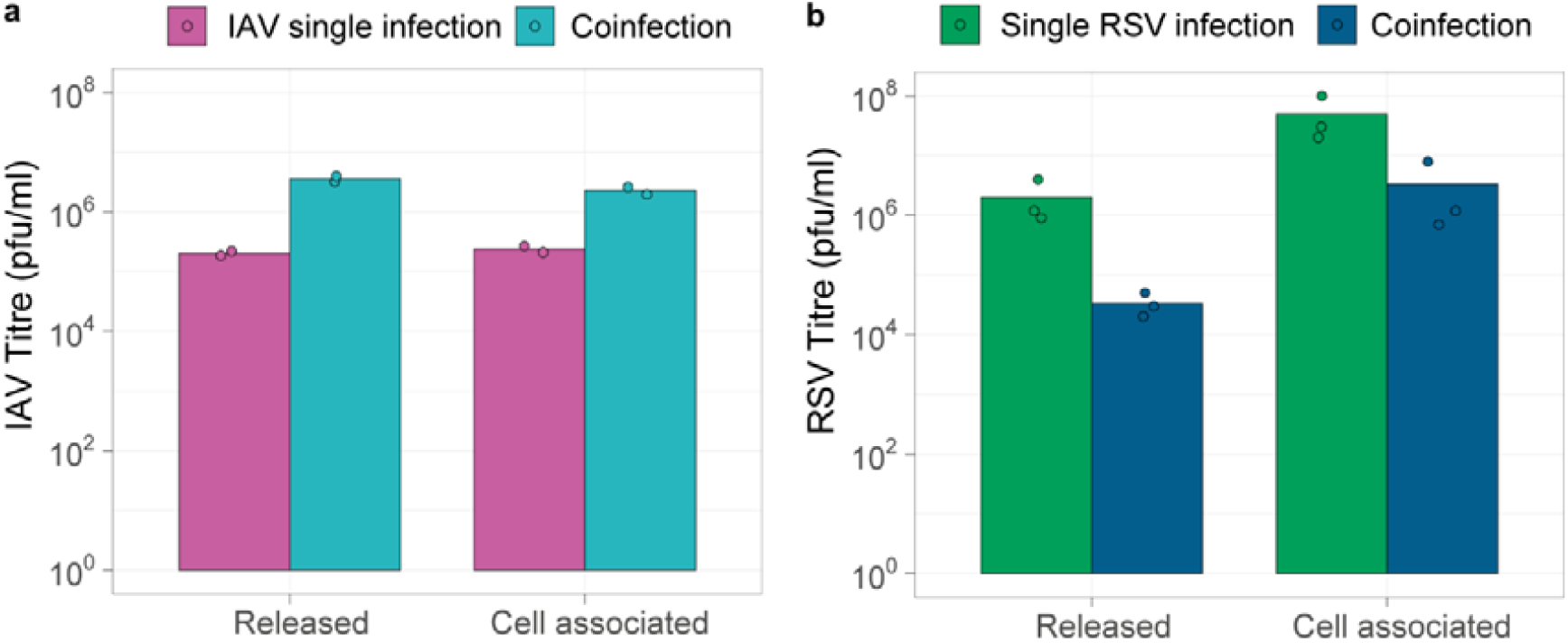
Supporting data for neutralisation assays. Quantification of viral input to neutralisation assays. Virus was harvested from supernatant (released) and cell associated fractions of single or coinfected A549 cells and transferred to neutralisation assays. The same virus stocks were then back titrated to determine infectious viral input. (a) Back titration of IAV in single infection (magenta bars) or coinfection (teal bars) by IAV plaque assay. (b) Back titration of RSV in single infection (green bars) or coinfection (dark blue bars) by RSV plaque assay.

**Extended Data Figure 7:**
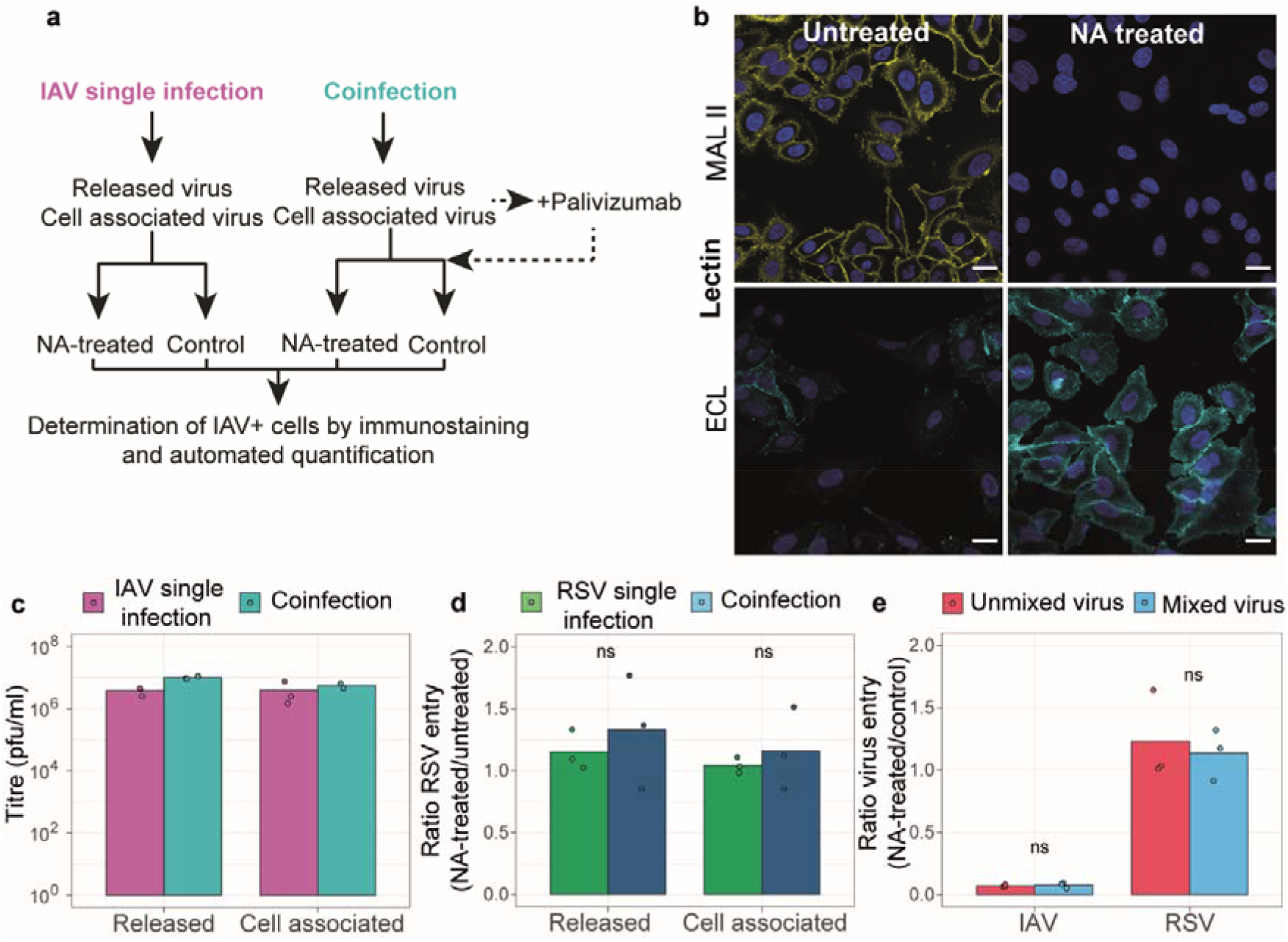
Supporting data for neuraminidase experiment. (a) Schematic demonstrating experimental design. (b) NA-treated and untreated cells stained with Maackia Amurensis Lectin II (MAL II) in yellow (top row) and Erythrina Cristagalli Lectin (ECL) in cyan (bottom row). (c) Viral input in pfu/ml of IAV as determined by back-titration of inoculum for NA-experiment by IAV plaque assay. (d) Ratio of RSV entry into NA-treated cells versus control cells when harvested from single infection (green bars) or mixed infection (blue bars). RSV entry to NA-treated cells was calculated as a percentage of the RSV-positive cell count in the matched untreated control. (e) Ratio of virus entry of IAV only or RSV only (red bars) into NA-treated over control cells, compared to entry of IAV pre-mixed with RSV or RSV pre-mixed with IAV into NA-treated over control cells (blue bars). Statistical significance was determined by Mann Whitney test, ns indicates p>0.05.

## Extended Data File

### Description of IAV and RSV particles

We identified structures consistent with IAV filaments and pleiomorphic particles that were densely coated with glycoproteins (Fig. 1A, Video S1). IAV filaments varied in length, with an average diameter of 84.8 (+/-6.97) nm. IAV ribonucleoproteins (RNPs) could be observed within pleomorphic virions (Figure 1A, Supplementary Video S1). We also identified RSV filaments (Figure 1B, Supplementary video S2) with an average width of 158 (+/-14.6) nm, but many irregular shaped virions were also observed (Fig. 2). Differences in glycoprotein distribution as well as RNP structure between RSV and IAV virions were evident. The glycoproteins in RSV virions exhibited sparser distribution along the virion than IAV glycoproteins (Figs. 1C and 1D). RNPs could be clearly seen in many IAV and RSV filaments. While IAV comprised a single ordered set of RNPs in a “7+1” arrangement at the distal end of filaments, multiple RNPs were detected within RSV filaments having herringbone morphology and ring-like structures attributed to be nucleocapsid (N) or N-RNA assemblies (*23, 52, 53*) (Fig. 1A-D, Fig. 2). Upon closer inspection of glycoproteins on the surface of virions (top and bottom slices in the tomograms), distinct patterns of glycoprotein arrangement were observed between the two viruses. In IAV virions, glycoproteins were densely packed, devoid of any long-range order, with the “triangular-shaped” trimeric head of HA clearly visible (Fig. 1E, 1F, Fig. 3). However, RSV exhibited a more symmetrical helical arrangement of glycoproteins (striped layers of density) with specific clustering in pairs (Fig. 1G, 1H, Fig. 3).

**Fig. 1.**
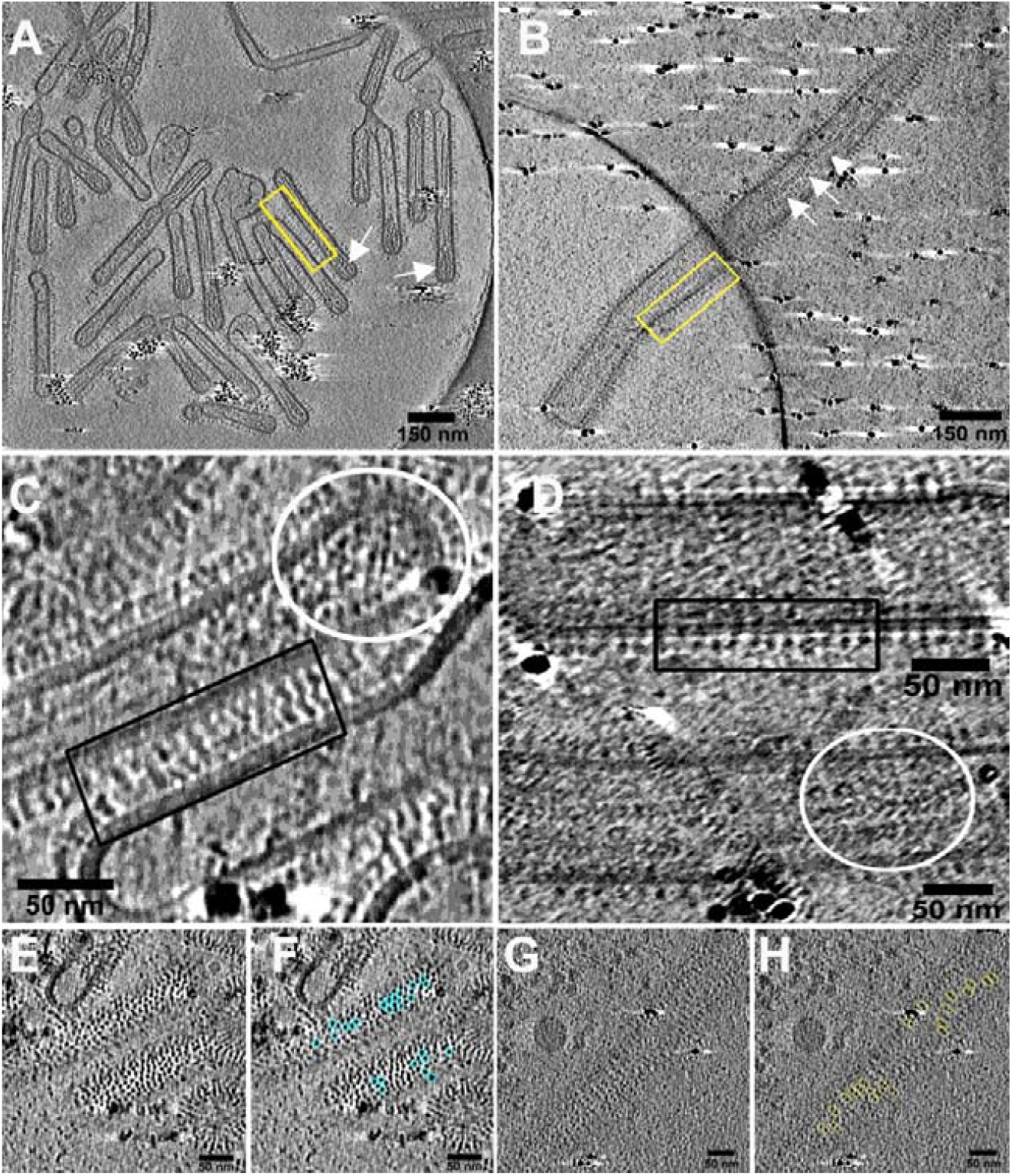
Cryo-ET shows differences in glycoprotein ordering and RNP organisation between IAV and RSV virions. Cryo-electron tomograms of IAV (**A**) and RSV **(B)** denote pleomorphic morphology alongside differences in glycoprotein and RNP organisation (yellow rectangle and white arrows). IAV virions exhibit a single set of ordered RNPs (**C**, white circle) while multi genomes can be seen in RSV virions (**D**, white circle). Glycoproteins can be seen in a picket fence like distribution along the viral membrane, although more densely packed in IAV than RSV (**C** and **D**, black rectangles). Looking at the glycoproteins from the top and bottom of the virions, significant differences are visible in glycoprotein packing: the triangular hemagglutinin HA spikes on IAV are densely packed with no long-range order (**E** and **F**, cyan circles) while RSV shows clustering of glycoprotein pairs (**G** and **H**, yellow ellipses).

**Figure 2.**
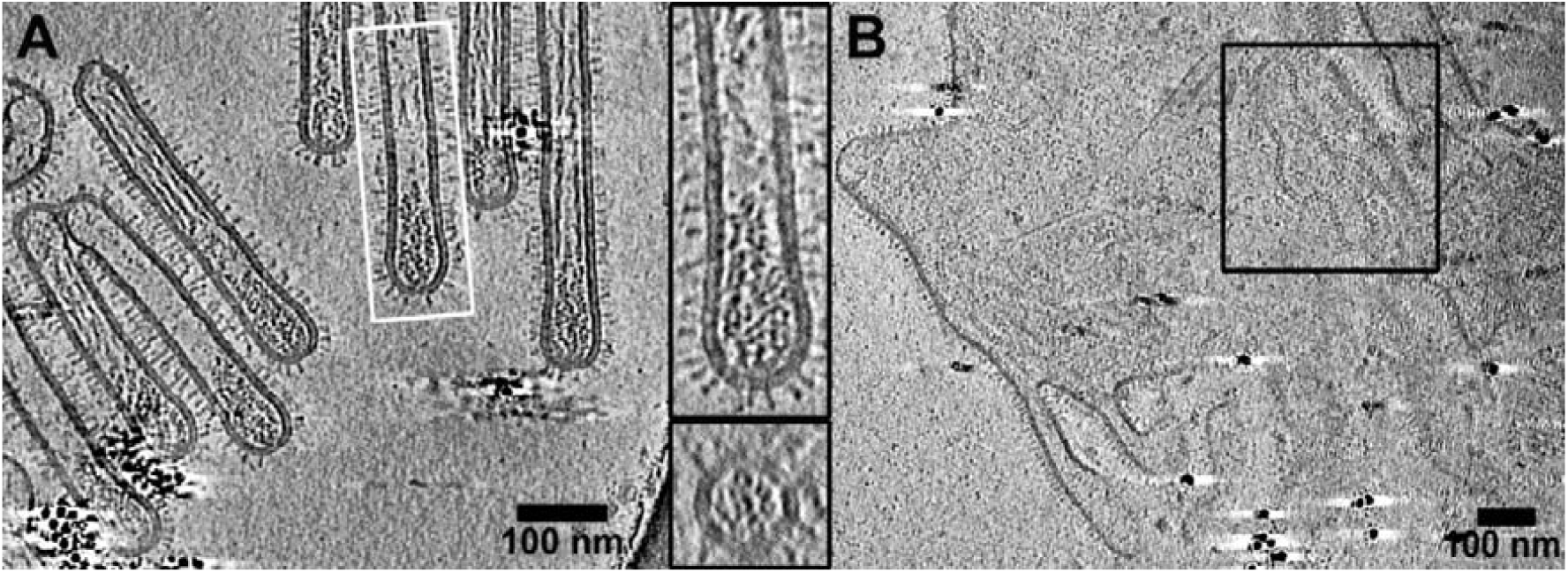
Distinct RNP organization in IAV and RSV virions seen by Cryo-ET. (**A**) IAV shows a single ordered set of RNPs at the distal end (white rectangle and magnified view of the same) of filaments. Transverse section (bottom image, magnified view of the white rectangle) of the filament denotes the “7+1” arrangement of RNPs in IAV virions. (**B**) Although RSV can form irregular shaped virions in addition to filaments, they are filled with multiple RNPs inside and decorated with spikes on the outside.

**Figure 3.**
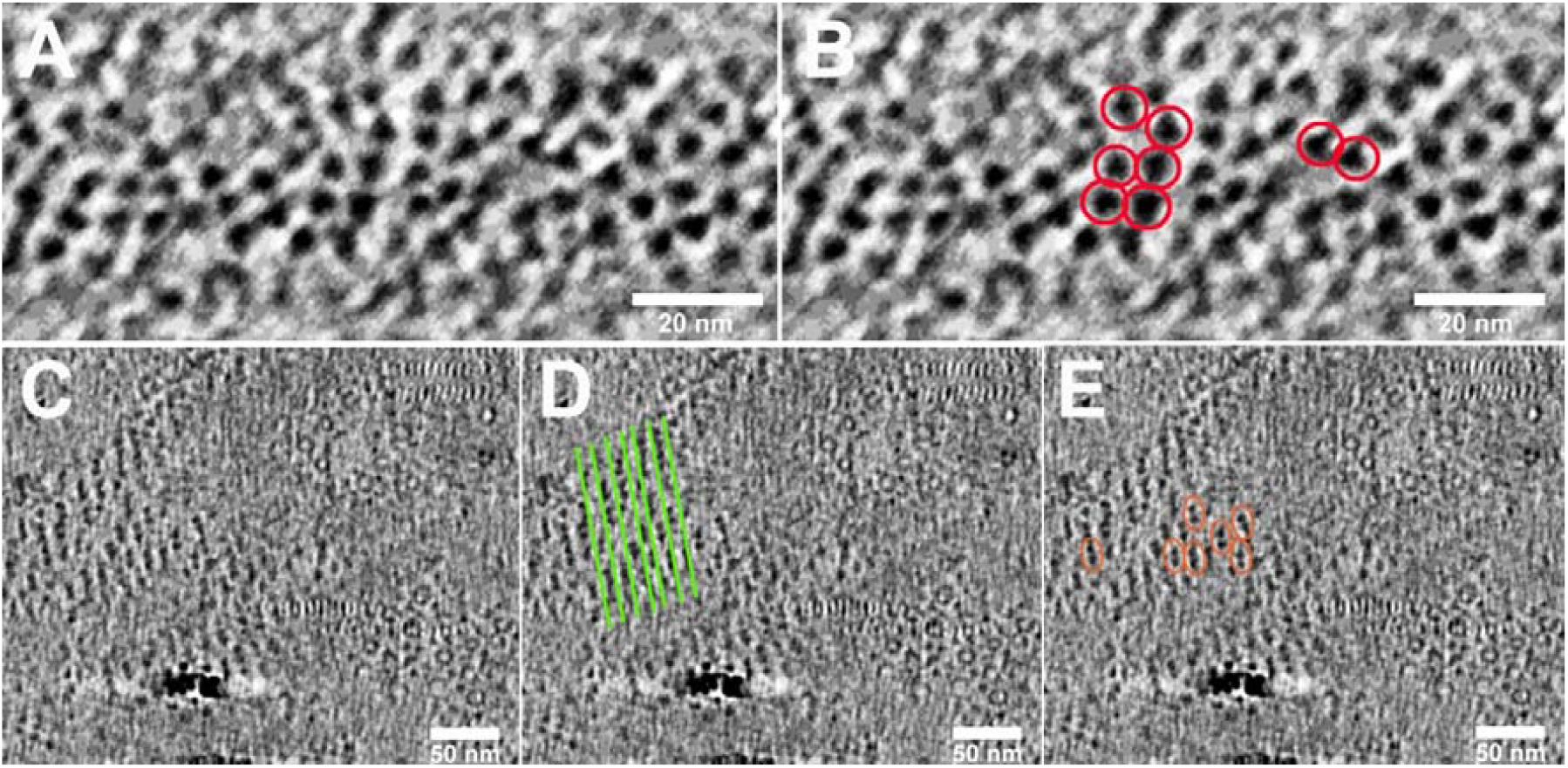
Different glycoprotein ordering between IAV and RSV virions. Inspection of top and bottom views of glycoprotein spikes (corresponding to top and bottom slices of the tomograms) on IAV and RSV virions indicates a different pattern of organization. (**A** and **B**) While the triangular heads (red circles) of the HA glycoprotein in IAV is clearly visible, the spikes are densely packed and devoid of any long-range order. (**C-E**) On the other hand the spikes on RSV appear to be helically ordered as seen by the striped arrays of density (**D**) and specific (**E**) clustering of glycoproteins in pairs (red ellipses) is clearly evident.

**Extended Data Movie 1**. Video showing serial sections through the z-axis of a tomogram of IAV virions exhibiting pleomorphic morphology. Glycoproteins and RNPs are labelled and denoted by arrows.

**Extended Data Movie 2**. Video showing serial sections through the z-axis of a tomogram of an RSV filament. Glycoproteins and RNPs are labelled and denoted by arrows.

